# Cell Surface-Bound La Protein Regulates The Cell Fusion Stage Of Osteoclastogenesis

**DOI:** 10.1101/2022.03.08.479741

**Authors:** Jarred M. Whitlock, Evgenia Leikina, Kamran Melikov, Luis Fernandez De Castro, Sandy Mattijssen, Richard J. Maraia, Michael T. Collins, Leonid V. Chernomordik

**Affiliations:** Section on Membrane Biology, Eunice Kennedy Shriver National Institute of Child Health and Human Development, National Institutes of Health, Bethesda, MD 20892, USA; Skeletal Disorders and Mineral Homeostasis Section, National Institute of Dental and Craniofacial Research, National Institutes of Health, Bethesda, MD, 20892, USA; Section on Molecular and Cell Biology, Eunice Kennedy Shriver National Institute of Child Health and Human Development, National Institutes of Health, Bethesda, MD 20892, USA

## Abstract

Multinucleated osteoclasts, essential for skeletal remodeling in health and diseases, are formed by fusion of osteoclast precursors with each fusion event raising their bone-resorbing activity. Here we report that nuclear RNA chaperone, La protein moonlights as an osteoclast fusion regulator. Monocyte-to-osteoclast differentiation starts with a drastic decrease in La levels. Then La reappears as a protease-cleaved species at the cell surface where it promotes fusion by mechanisms independent of La-RNA interactions. Appearance-and-disappearance of cell-surface La act as an on-and-off switch of the fusion activity with fusion slowing down when surface La is replaced in mature osteoclasts by full-length nuclear La. Inhibiting surface La in a novel explant model of fibrous dysplasia inhibits excessive osteoclast formation characteristic of this disease, highlighting La’s potential as a therapeutic target.

**One-Sentence Summary:** A nuclear RNA chaperon moves to the surface of osteoclasts to control their formation and function in bone metabolism.

## INTRODUCTION

Bone-resorbing osteoclasts are responsible for essential, life-long skeletal remodeling, and their dysfunction is a major contributor to bone diseases affecting >200 million individuals worldwide ^1^, including osteoporosis, fibrous dysplasia (FD), Paget’s disease and osteopetrosis ^2–6^. Multinucleated osteoclasts are formed by the successive fusion of mononucleated precursor cells ^7^. The number of nuclei per syncytial osteoclast, thus, the number of fusion events that generated each cell, directly correlates with the cell’s ability to resorb bone ^8–10^, as reflected in the fact that the number and size of osteoclasts are significantly altered in many bone diseases ^11,12^. Recent studies suggest that during their relatively long lifetime ^13^ osteoclasts can go through additional rounds of cell fusion. Following their initial formation, multinucleated osteoclasts can undergo fission producing smaller daughter cells termed osteomorphs that can then migrate and fuse again to form mature multinucleated osteoclasts in a different location ^14^. Despite the fundamental role of cell-cell fusion in osteoclast formation and bone remodeling, the mechanisms underpinning this process as well as other cell-cell fusion processes in normal physiology and in diseases ^15–17^ remain to be fully understood. A number of proteins, including DC-STAMP, OC-STAMP, syncytin 1, annexin A5 (Anx A5), S100A4, CD47 and SNX10 ^18–22^, have been implicated in osteoclast fusion, however, how osteoclasts regulate their fusion and arrive at the “right size” to fulfil their biological function remains elusive ^23^.

Osteoclasts derive from monocytes when stimulated by macrophage colony stimulating factor (M-CSF), receptor activator of NF-kappaB ligand (RANKL), and other cytokines released by bone-forming osteoblasts and osteocytes ^24^. In *vitro*, M-CSF and RANKL together are sufficient to elicit osteoclastogenesis. First, M-CSF stimulates generation of adherent mononucleated osteoclast precursors. Second, RANKL commits these precursors to osteoclastogenesis and fusion ^25^. While exploring proteomic changes during this stepwise process, we unexpectedly discovered that osteoclastogenesis involves lupus La protein (*SSB* gene product). La, also referred to as LARP3 and La autoantigen, is generally recognized as an abundant and ubiquitous RNA-binding protein ^26^. La has a nuclear localization sequence (NLS) at its C-terminus as well as multiple other intracellular trafficking signals ^27^ that result in La being observed almost exclusively in the nucleus of human cells ^28^. The best characterized function of nuclear La is to protect precursor tRNAs from exonuclease digestion through specific interactions between La’s highly conserved, N-terminal La domain and the 3’ ends of tRNA. In addition to its nuclear functions, La shuttles to the cytoplasm ^29^ and assists in the correct folding of mRNAs, acting as an RNA chaperone, where both the N- and C-terminal halves of human La appear capable of RNA strand annealing and dissociation ^30^. In a few specialized biological processes (e.g., apoptosis, viral infection, serum starvation), La protein is dephosphorylated at phospo-Ser-366, loses its NLS via proteolytic cleavage, and this low molecular weight (LMW) species trafficks to the surface of the cells ^27,31–34^. However, the biological function of this cleaved, surface La, if any, is unknown.

Here, we report that osteoclast formation is accompanied by and depends on drastic changes in the steady-state level, molecular species, and intracellular localization of La protein. We demonstrate that La functions as a regulator of osteoclast fusion and impacts osteoclasts’ ability to resorb bone. Surprisingly, La, present in primary human monocytes, nearly disappears in M-CSF derived osteoclast precursors. RANKL-induced commitment to osteoclastogenesis drives the reappearance of La protein in a cleaved form at the surface of committed, fusing osteoclasts. As osteoclast fusion plateaus, cleaved La disappears and higher molecular weight, full-length protein (FL-La) is observed within the nuclei of mature, multinucleated osteoclasts. Perturbing La expression, cleavage or surface function inhibits osteoclast fusion, while exogenous, surface La promotes fusion. Moreover, the mechanism by which La promotes osteoclast fusion is independent of La’s ability to interact with RNA through its highly conserved La domain. Indeed, a C-terminal portion of La, lacking the La domain and RNA recognition motif 1 (RRM1) is sufficient to promote fusion between human osteoclasts. Our findings indicate that La protein, which plays an ancient, well-described and essential role in the RNA biology of eukaryotes, has been adapted in mammals to also serve as an osteoclast fusion manager. In this new, highly specific role on the surface of fusing osteoclasts, La may present a promising target for treatment of bone diseases stemming from perturbed bone turnover.

## RESULTS

### Formation of multinucleated osteoclasts involves La protein

Human osteoclastogenesis was modeled by treating primary monocytes with M-CSF to derive mononucleated osteoclast precursors to which recombinant RANKL was later added to obtain multinucleated osteoclasts that readily resorb bone ^18^ (Fig. 1a,b, Fig. S1a-c). Osteoclast precursors begin fusing at ~2 days following RANKL addition and after ~5 days reach the sizes (~ 5-10 nuclei/cell) characteristic of mature multinucleated osteoclasts ^10,35,36^ (Fig. 1b, Fig. S1c).

**Figure 1:**
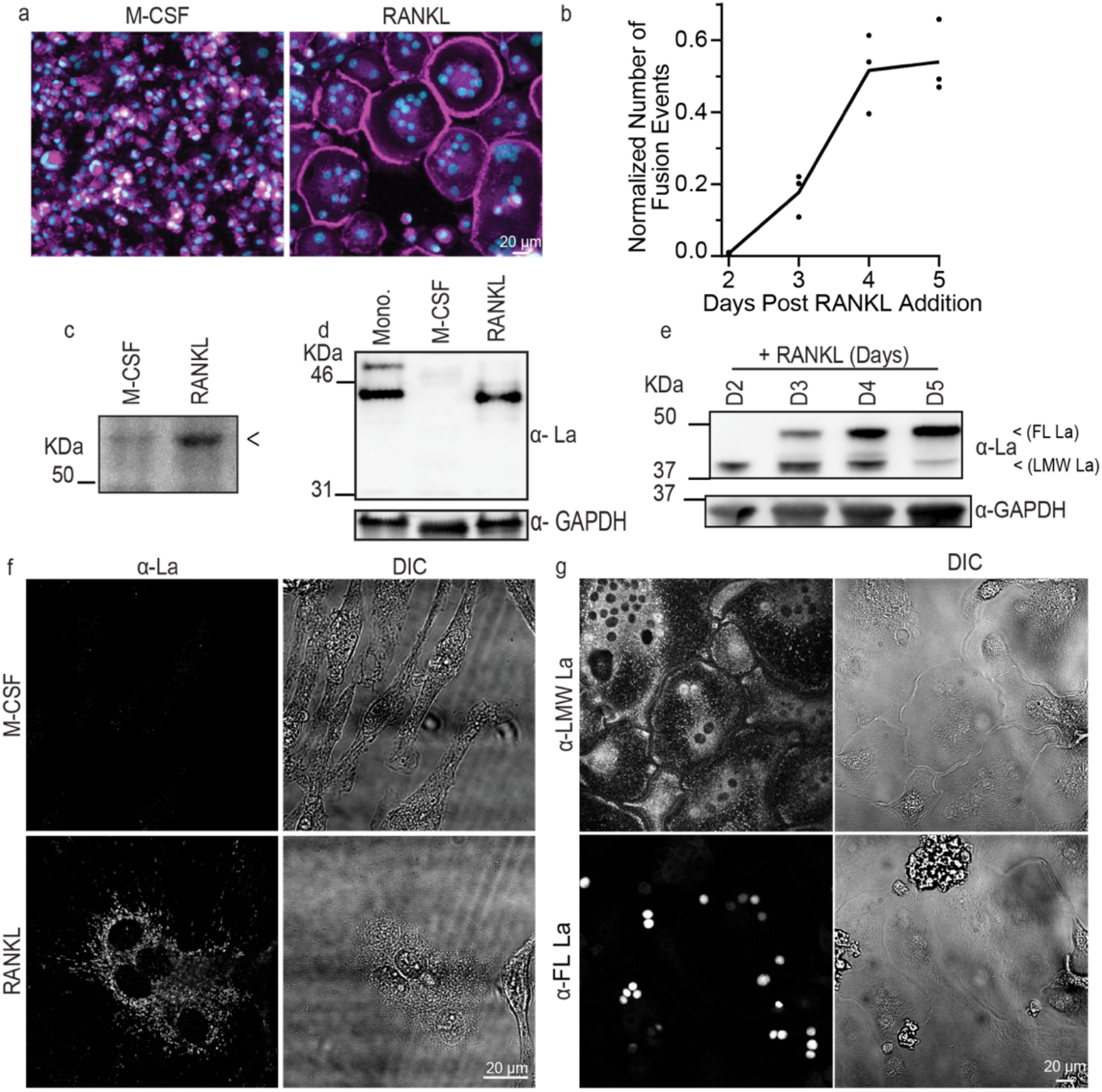
Osteoclastogenic differentiation is accompanied by drastic changes in the steady-state levels and localization of La molecular species. **(a)** Representative images of stages of osteoclastogenic derivation of human monocytes following M-CSF (6 days) and M-CSF + RANKL (5 days), respectively. (Magenta = Phalloidin-Alexa488, Cyan = Hoechst) **(b)** Quantification of the number of fusion events normalized to the total number of nuclei observed over time following RANKL addition. (n=3) Each point represents an average of >7,500 nuclei scored. **(c)** Representative Bis-Tris PAGE separation and silver staining of whole protein lysates from the osteoclastogenic stages depicted in **a**. Lysates were ran until the 50 KDa marker nearly ran off the 4-12% polyacrylamide gel to achieve maximal separation of proteins at this molecular weight, leading to the band of interest appearing misleadingly heavy. < denotes band of interest excised from both lanes and evaluated via mass spectrometry. **(d)** Representative Tris-Glycine Western blot evaluating La expression in whole protein lysates from primary human monocytes and the osteoclastogenic stages depicted in **a**. (α-La, Abcam 75927) **(e)** Representative tris-glycine Western blot evaluating the time course of La expression following RANKL addition. (α-La, Abcam 75927) (α-GAPDH loading control) **(f)** Representative immunofluorescence images of La in the M-CSF derived osteoclast precursors and at 3 days post RANKL application (α-La, Abcam) **(g)** Representative immunofluorescence images of La in osteoclasts using La antibodies specifically recognizing LMW La (α-LMW La) and FL La (α-FL La). Cells were fixed 4 days post RANKL addition, when both LMW and FL La are abundant.

While evaluating proteomic changes associated with osteoclastogenesis, we discovered a distinct protein that was nearly absent in M-CSF derived precursors but abundantly expressed in osteoclasts following ~3 days of RANKL stimulated osteoclastogenesis, when the cells were rapidly fusing (Fig. 1c, arrow). Using mass spectrometry analysis, we identified this protein as La (Fig. S1d). The low level of La in M-CSF-derived macrophages was unexpected, as La is generally considered an abundant, ubiquitous protein ^26,37–40^.

Western blot analysis confirmed that La is well expressed in monocytes, markedly reduced in M-CSF-derived osteoclast precursors and La’s high steady-state levels return during RANKL-induced osteoclast formation (Fig. 1d). Our data suggest that La’s tight regulation during osteoclastogenesis is likely carried postranslationally, as M-CSF derived precursors contain even more La transcript (gene SSB) than after RANKL application (Fig. S1e). La appears as two distinct, temporally separated molecular species during osteoclastogenesis (Fig. 1e). A low molecular weight (LMW La) species detected at the timepoints at the onset and during robust fusion is replaced by a higher molecular weight species, corresponding to fulllength La (FL La), as fusion slows and osteoclasts reach a mature size.

In addition to changes in molecular weight, osteoclastogenic differentiation of human monocytes is accompanied by a dramatic change in La’s location within cells. Canonically, La exhibits robust nuclear staining ^26^, as illustrated for HeLa cells (Fig. S1f). In contrast, M-CSF derived osteoclast precursors exibit minimal La staining (Fig. 1f), consistent with our biochemical analysis (Fig. 1d). Addition of RANKL produced abundant La signal in committed, fusing osteoclasts, however, in contrast to other human cell types and tissues ^26,27^, La appears as distinct, predominantly non-nuclear puncta throughout fusogenic osteoclasts during early stages of osteoclast fusion (Fig. 1f).

The anti-La antibody (a-La) used in Fig. 1d recognizes both LMW La and FL La species. To determine whether the different molecular weight species of La exhibit different localization during osteoclastogenesis, we analyzed several commercially available antibodies and selected antibodies that preferintally recognized LMW La species (a-LMW La, Methods and Table S1 and Fig. S1g). FL La is largely phosphorylated at Ser366 and localizes to the nucleus ^41^. Previous work has demonstrated that LMW La is produced by the cleavage of FL La, but that FL La must be dephosphorylated at 366 before FL La can be cleaved ^33^. Moreover, previous reports have shown that antibodies specific for phosphoSer366 La do not recognize LMW La ^33^. So, we utilized α-phosphoSer366 La antibodies to preferintally stain for FL La (α-FL La). At intermediate timepoints, where both La molecular species were present, we found that LMW La was predominantly detected outside the nucleus, throughout the cell, while FL La was observed exclusively in the nuclei of fused cells (Fig. 1g). We further confirmed the shift in La distribution during osteoclastogenesis by staining with a-La at different days post RANKL application (Fig. S1h).

Finding that osteoclastogenic differentiation is accompanied by drastic changes in the expression and localization of La motivated us to explore whether La is functionally involved in osteoclast formation. We found that La expression tightly regulates the formation of multinucleated osteoclasts. RNAi mediated reduction of La transcript *(SSB)* drastically inhibited osteoclast fusion (Fig. 2a-c). Cytosolic localization of La at the time of fusion was also observed during osteoclastogenic differentiation and fusion of RAW 264.7 derived murine osteoclast precursors (Fig. 2d). Furthermore, Western blot analysis of cell lysates collected separately from mostly mononucleated cells and from mostly multinucleated cells (see Methods) indicated that the robust fusion at day 3 post-RANKL was accompanied by a drastic increase in steady-state levels of La (Fig. 2e). These findings suggested that La dependence in osteoclast formation is conserved in humans and mice. Note that in distinction to human cells, in the case of RAW 264.7 cells, following the stage of active fusion by day 5, the levels of La returned to lower pre-fusion levels.

**Figure 2:**
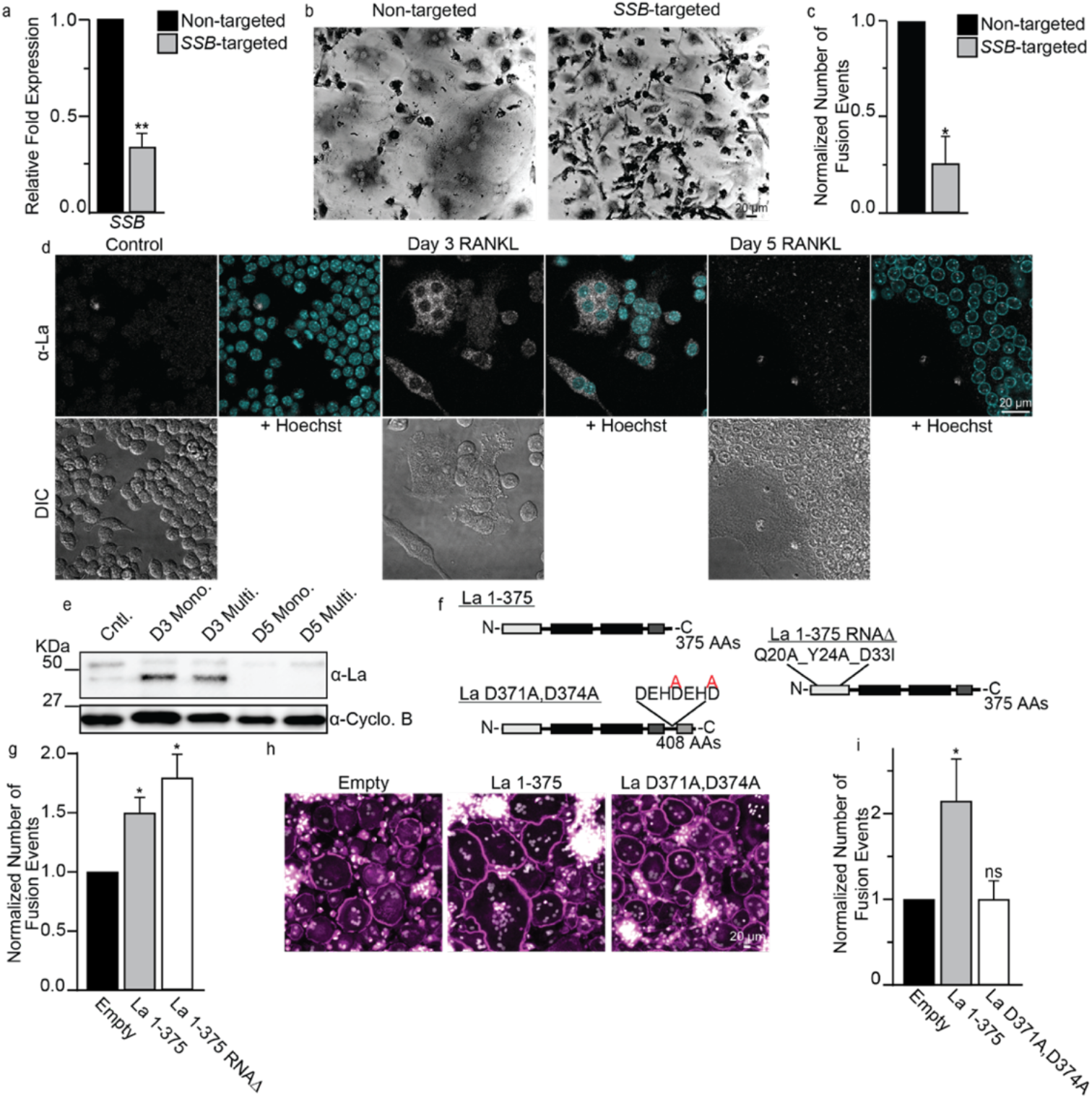
Osteoclast formation depends on truncated La, but the function of the La domain is dispensable. **(a)** qPCR evaluation of *SSB* following siRNA treatment of human osteoclast precursors. (n=4) (P=0.0043) **(b)** Representative, phase contrast images of non-targeted and *SSB*-targeted human osteoclasts, as in **a**, stained for TRAP. **(c)** Quantification of the number of fusion events in formation of syncytia with 3+ nuclei in the experiments like the one in **b**. (n=3) (P=0.0306) **(d)** Representative immunofluorescence images of La in RAW 264.7 prior to mRANKL addition (Cntl.), 3 days post mRANKL addition and 5 days post mRANKL addition when we routinely observe massive, multinucleated osteoclasts like the one imaged here. **(e)** Representative tris-glycine Western blot of whole cell lysates taken from murine, RAW 264.7 treated as in **d**. mRANKL treated cells were enriched into mononucleated (Mono.) or multinucleated (Multi.) populations as described in the Methods. · (Cyclophilin B (-Cyclo B) loading control) **(f)** Topological illustrations of LMW La 1-375, “uncleavable” La D371A,D374A and “RNAΔ” La 1-375 Q20A_Y24A_D33I. **(g)** Quantification of the number of fusion events in syncytia with 3+ nuclei in RAW 264.7 cells transfected with empty, La 1-375 or La 1-375 Q20A_Y24A_D33I expression plasmids. (n=3) (P=0.0213 and 0.0173, respectively). **(h)** Representative fluorescence images of human monocyte-derived osteoclasts transfected with empty, La 1-375 or La D371A,D374A expression plasmids. (Magenta = Phalloidin-Alexa488, Grey = Hoechst) **(i)** Quantification of the number of fusion events in syncytia with 3+ nuclei in **h**. (n = 4) (P=0.0205 and 0.325, respectively). Statistical significance was evaluated via paired t-tests. * = P<0.05. ** = P<0.001. Error bars = SEM.

To identify the functional form of La associated with osteoclastogenesis, we focused on the relationship between the appearance of LMW La and fusion between osteoclasts. Earlier reports demonstrate that during apoptotic progression human La is cleaved by caspases at Glu-375, removing La’s NLS ^33,42^. We found that overexpression of La 1-375, mimicking this cleaved species, greatly promoted fusion in both RAW 264.7 derived, murine osteoclasts and monocyte derived, human osteoclasts (Fig. 2f-i). In contrast, an uncleavable mutant of FL La, D371A,D374A La (point mutations disrupting La’s predicted caspase cleavage sites) had no effect on osteoclast fusion despite being expressed at similar levels, suggesting that formation of multinucleated osteoclasts depends on LMW La (Fig. 2h,i and Fig. S2). Furthering this point, we found that the pan caspase inhibitor z-VAD blocked the production of LMW La in differentiating osteoclasts (Fig. S3a). Blocking the caspase-dependent production of LMW La resulted in the premature retention of FL La within the nuclei of unfused osteoclasts (Fig. S3c vs b) and significantly perturbed the ability of osteoclasts to form multinucleated syncytia (Fig. S3d, see also ^43^). Taken together our data support the role of caspase-cleaved, LMW La in the formation of multinucleated osteoclasts.

To summarize, osteoclastogenesis is accompanied by drastic changes in La levels, molecular species and location within fusing osteoclasts. A cleaved, non-nuclear La species promotes osteoclast formation, and as cells arrive at a mature size, LMW La is replaced by FL La detected in the nuclei of syncytial osteoclasts.

### Cell surface associated La regulates the cell fusion stage of osteoclast formation

In our characterization of La’s role in osteoclastogenesis, we first explored whether La exerted its function in the formation of osteoclasts indirectly by altering the expression of factors implicated in osteoclastogenic differentiation or osteoclast fusion. While La expression in many cell types influences the steady-state levels of many transcripts/proteins ^39^, RNAi suppression of La did not alter the steadystate transcript levels of the essential osteoclastogenesis factors NFATc1 and CTSK or transcripts coding for the fusion associated proteins syncytin 1, Anx A5, S100A4 or the lipid scramblase anoctamin 6/TMEM16F (Fig. S4) ^18^. Therefore, while La knockdown inhibited the formation of osteoclast syncytia, it did not grossly impact osteoclast differentiation or other machinery critical for cell-cell fusion. To further explore the mechanism by which La influenced the formation of osteoclasts but not their differentiation, we assessed whether the formation of multinucleated osteoclasts depends on La’s well characterized RNA binding function. This highly conserved function is based on high affinity interactions between the La domain and its high affinity oligo(U)-3’ binding site common to RNA polymerase III transcripts. To assess the requirement of high affinity interactions between the La domain and transcripts in osteoclastogenesis, we overexpressed a mutant La 1-375 with three-point mutations known to functionally impair La domain function, Q20A/Y24A/D33I ^44,45^ (La 1-375 RNAD). We found that La 1-375 RNAD promoted formation of multinucleated osteoclasts as robustly as wild type La 1-375, indicating that the La domain’s high-affinity for RNA polymerase III transcripts is dispensable for La’s role in osteoclast formation (Fig. 2f,g).

Our second piece of evidence suggesting that La’s role in the formation of multinucleated osteoclasts was not through La’s canonical role in RNA metabolism was the demonstration that La’s function in osteoclasts was not in the nucleus or the cytoplasm, but rather, at the cell surface. As noted earlier, in differentiating osteoclasts La loses its NLS and appears in punctate structures throughout the cell. We enriched proteins from RANKL committed osteoclasts at timepoints when cells were actively fusing into soluble, cytosolic or membrane-associated protein fractions. As expected, we found actin mostly in the cytosolic fraction, transmembrane RANK receptor in the membrane fraction, and the peripheral membrane protein Anx A5 in both fractions (Fig. 3a). While La protein is putatively considered a soluble protein, in differentiating osteoclasts, we found La in both cytosolic and membrane-associated fractions, suggesting La was unexpectedly associating with membranes during osteoclastogenesis (Fig. 3a).

**Figure 3:**
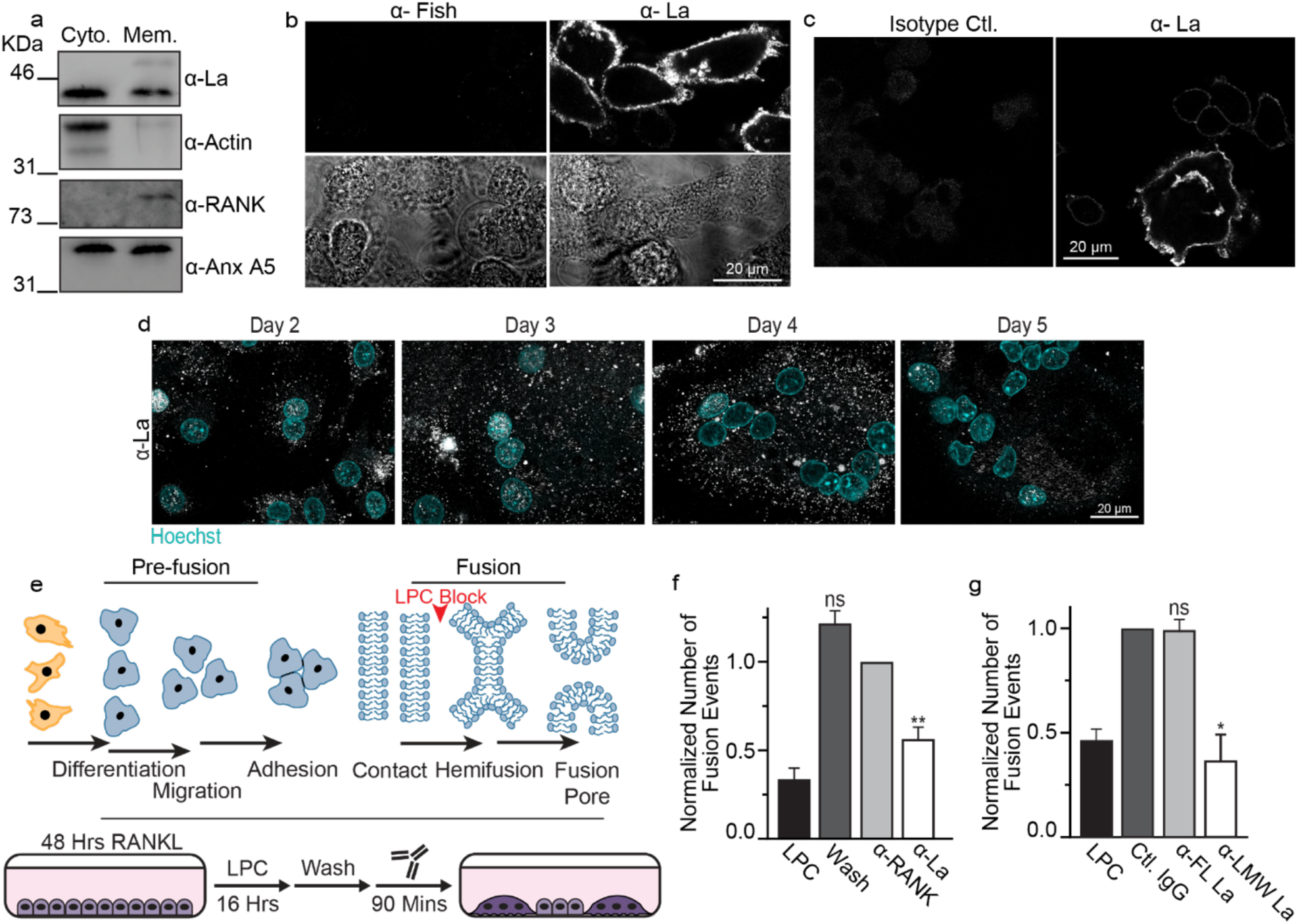
La associates with membranes, traffics to the surface and controls osteoclast membrane fusion. **(a)** Westerns of cytosolic vs membrane associated protein fractions from human osteoclasts. **(b)** Representative immunofluorescence images comparing surface staining of α-Fish/TKS5 or α-La antibodies in human osteoclasts under non-permeabilized conditions (top) and DIC (bottom). **(c)** Representative immunofluorescence images comparing surface staining of isotype control or α-La antibodies in RAW 264.7 derived osteoclasts under non-permeabilized conditions. **(d)** Representative immunofluorescence images of La in forming osteoclasts 2-5 days post RANKL application (α-La, Abcam). Cells were stained for La at the described timepoints without membrane permeabilization (lower panel). **(e)** Cartoons illustrating the stepwise process of the formation of multinucleated osteoclasts (top), and our approach for isolating membrane fusion stage from the preceding stages of osteoclast differentiation (bottom). Application of the hemifusion inhibitor LPC following 48 hours of RANKL elicited osteoclastogenesis allows pre-fusion differentiation stages but blocks hemifusion, synchronizing cells. Removing LPC allows us to specifically probe membrane fusion between osteoclasts. **(f)** Quantification of human osteoclast fusion decoupled from pre-fusion stages and synchronized as depicted in **e** with fusion in the presence of a-La and with no antibodies added (Wash) normalized to α-RANK control. (n=3) (P=0.0086 and 0.1330, respectively) **(g)** The effects of a-FL La or a-LMW La application on synchronized fusion of human osteoclasts. Data normalized to those after application of isotype control (IgG). (n=2) (P=0.03 and 0.4581, respectively). (**e, f**) “LPC” – fusion observed without removal of LPC. Statistical significance was assessed via paired t-test. * = P<0.05. ** = P<0.001. Error bars = SEM.

In earlier reports, La cleavage in apoptotic cells was associated with the detection of La on the cell surface ^33,42^, however whether this surface La played some cellular function or operated simply as a surface antigen remains unknown. To assess whether osteoclast La traffics to the cell surface following cleavage, we stained fusing osteoclasts with α-La antibodies under non-permeabilizing conditions (Fig. 3b,c). In contrast to the osteoclast peripheral membrane protein Fish ^46^, which is enriched during osteoclast fusion and binds to the cytoplasmic leaflet of the plasma membrane (PM) ^46^, La abundantly decorated the surface of fusing human osteoclasts (Fig. 3b). Moreover, this surface pool of La is not exclusive to human osteoclasts. We also observed significant La surface staining in RAW 264.7 derived, murine osteoclasts (Fig. 3c), suggesting surface La is a feature common to fusing osteoclasts in mammals.

Using surface staining with a-La at different days post RANKL application, we observed the transient increase in La at the time ponts associated with robust fusion (Fig. 3d vs. Fig. 1b), further implicating La in fusion.

We then assessed whether La at the surface of human osteoclasts functions at the cell fusion stage of osteoclastogensis. All cell-cell fusion events in development and tissue maintenance proceed through slow (days), asynchronous differentiation processes that prepare fusion competent cells ^15^. Then, PM fusion occurs by the rapid (minutes) progression from the initial formation of hemifusion connections to fusion pores that unite the volumes of two cells (Fig. 3d, top). We decouple these steps in the formation of multinucleated syncytia using the hemifusion inhibitor lysophosphatidylcholine (LPC) ^18^. LPC’s inverted cone shape is not conducive to the concave geometry of the hemifusion stalk, so ready-to-fuse cells are trapped upstream of hemifusion. After removing LPC, cells undergo synchronized fusion relatively rapidly (within 90 mins), affording us the ability to assess the function of proteins specifically in the membrane fusion stage of osteoclast formation decoupled from upstream differentiation processes (Fig. 3d, bottom). We accumulated ready-to-fuse, RANKL committed cells in the presence of LPC, and then lifted this hemifusion blockade by washing out LPC. Application of a-La antibodies at the time of LPC removal significantly inhibited synchronized, osteoclast membrane fusion (Fig. 3e). In contrast, antibodies targeting the PM receptor RANK at the surface of osteoclasts had no effect (Fig. 3e). While RANKL-RANK signaling triggers upstream osteoclastogenic differentiation, inhibition of RANK following hemifusion synchronization fails to inhibit membrane fusion, as fusion itself does not depend on the activity of RANK ^18^. Moreover, while a-LMW La antibodies completely blocked synchronized osteoclast membrane fusion, a-FL La antibodies had no effect (Fig. 3f).

In contrast to the fusion-inhibiting effects of antibodies targeting surface La, application of recombinant La dramatically promoted osteoclast fusion. Application of FL La (La 1-408), truncated La (La 1-375) or truncated, RNA binding mutant La 1-375 RNAΔ outside fusing osteoclasts significantly promoted the formation of multinucleated syncytia (Fig. 4a-c). This promotion was not observed when recombinant La was heat inactivated. Recombinant La 1-375 RNAΔ promoted fusion similarly to La 1-408 and La 1-375, confirming that La’s high-affinity interactions with RNA polymerase III transcripts are not required for La’s role in regulating osteoclast fusion (Fig. 4a-c). Moreover, the ability of FL La to promote osteoclast fusion demonstrates that FL La itself is not fusion incompetent, but rather suggests that proteolytic processing and/or dephosphorylation of La are important for its delivery to the cell surface.

**Figure 4:**
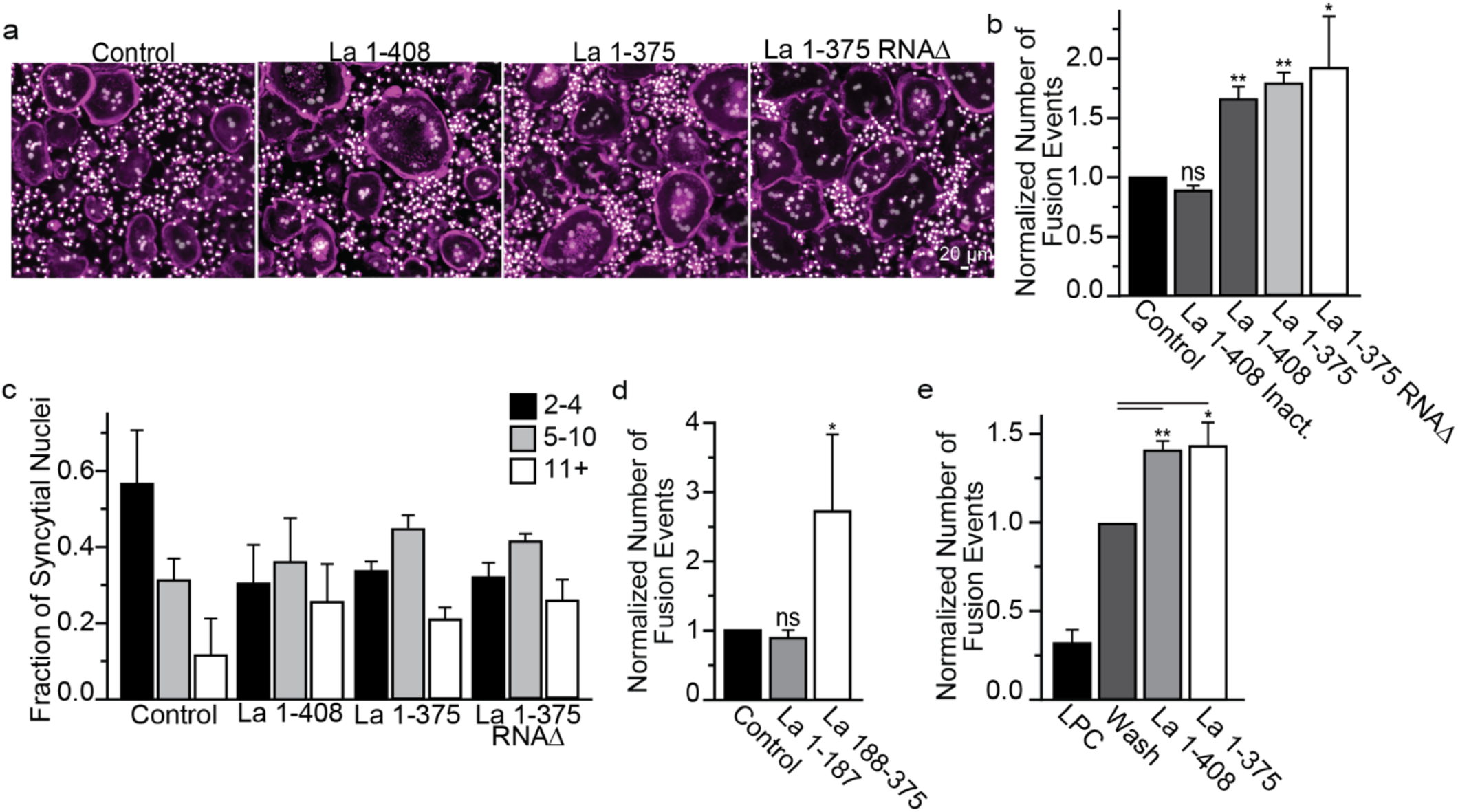
Recombinant La promotes osteoclast fusion. **(a)** Representative fluorescence images of human osteoclasts 3 days post RANKL addition without or with the overnight (end of day 2 post RANKL) addition of recombinant heat-inactivated La 1-408, La 1-408, La 1-375 or La 1-375 Q20A/Y24A/D33I. Recombinant proteins were added at ~40 nM at the end day 2 post-RANKL addition, and cells were fixed the next morning. (Magenta = Phalloidin-Alexa488, Grey = Hoechst) **(b)** Quantification of **a.** (inactivated n=2; others n=3,) (P=0.1232, 0.0015, 0.0035 and 0.0491, respectively) **(c)** Quantification of the fraction of nuclei in fused cells that were present in syncytia of various sizes from **a**. (n=3). **(d)** Quantification of the number of fusion events with or without the addition of La 1-187 or La 188-375. Recombinant proteins were added at ~40 nM at the end day 2 post-RANKL addition, and cells were fixed the next morning. **(e)** The quantification of synchronized fusion events (as illustrated in Figure 3d) without (wash) and with addition of recombinant La species. “LPC” – indicates that the hemifusion inhibitor was left until fixation. (n=3) (P=0.001 and 0.03, respectively.) Statistical significance was evaluated via paired t-test. In (b, d, e) the data were normalized to those in control (no protein added in b, d, and wash with no proteins added e). Error bars = SEM.

To further resolve the contributions of La domains critical for RNA binding, La and RRM1 domains ^26,27,44,45^, we split La 1-375 into La 1-187 and La 188-375. We found that La 188-375 greatly promoted the formation of multinucleated osteoclasts, whereas La 1-187 had no effect (Fig. 4d). These data demonstrate that the La domain, RRM1 and La’s C-terminal 33 AAs are dispensable for La’s role in osteoclast formation (Fig. 4d). Importantly, La promotes the formation of osteoclasts at the membrane fusion stage rather than some pre-fusion stage of the differentiation. To this point, application of recombinant La to LPC-synchronized osteoclasts dramatically promotes osteoclast membrane fusion (Fig. 4e).

All these data indicated that La functions at the cell surface during the membrane fusion stage of osteoclast formation to promote the formation of large, multinucleated osteoclasts. Proteins involved in membrane fusion can be divided into protein fusogens that are sufficient for generating hemifusion intermediates and opening of fusion pores, and proteins that regulate fusogen activity ^15^. To test whether cell surface La may fuse membranes on its own, functioning as an active protein fusogen, we assessed La’s ability to promote fusion between 3T3 fibroblasts, stably expressing HA0 (an uncleaved form of the influenza fusogen hemagglutinin (HA)), and red blood cells (RBCs) labeled with lipid and a content probes ^47^. HA0 is fusion-incompetent but establishes very tight contacts between HA0-expressing fibroblasts and RBCs. We found that none of 872 analyzed HA0-cell bound RBCs exchanged lipid (hemifusion indicator) or cytoplasmic (fusion pore indicator) probes in response to application of 40 nM recombinant La. Based on Wilson’s method ^48^, the probability of La mediated fibroblast-RBC fusion does not exceed 0.0044 per cell contact and, under these conditions, exhibits at least 100-fold lower activity than the fusion mediated by a *bona fide* fusogen - activated HA (~0.5 per contact). These data indicate that La does not itself have detectable cell-cell fusogenic activity, but rather, La likely regulates some larger fusion mechanism specific to osteoclasts.

The lack of a direct fusogenic activity of La suggested that cell surface La interacts with some other protein(s) involved in fusion. To test this hypothesis, we assessed whether La interacted with Anx A5, a peripheral membrane protein, also involved in membrane fusion stage of osteoclast formation (^18,49^) and upregulated at similar timepoints in osteoclastogenesis ^18^. We immunoprecipitated La and La-containing protein complexes from fusing human osteoclasts on magnetic beads with a-La Abs and found that La protein complexes contained Anx A5 (Fig. S5a). La supramolecular complexes from fusing osteoclasts contained neither Anx A1 nor Anx A4, both abundant in fusing osteoclasts (Fig. S5b), demonstrating specificity in La’s association with Anx A5. Moreover, immunoprecipitations of Anx A5 supramolecular complexes from fusing human osteoclasts isolated using a-Anx A5 Abs, contain La (Fig. S5b).

Additional evidence for La-Anx A5 interactions came from the experiments in which we found that direct binding of La to Anx A5 anchors La on phosphatidylserine (PS) containing phospholipid liposomes. Anx A5 binds to PS containing membranes in a Ca^2+^-dependent manner. We introduced recombinant La alone or along with recombinant Anx A5 to PS containing liposomes and pelleted liposomes by centrifugation to evaluate whether La was enriched with liposomes or in supernatant (Fig. S5c). La alone did not pellet along with liposomes. In contrast, both La and Anx A5 pelleted with liposomes in response to Ca^2+^ (Fig. S5d). La membrane association required Anx A5, Ca^2+^ and PS, as neither La nor Anx A5 pelleted with liposomes lacking PS (Fig. S5d). Our findings suggest that direct interactions between La and extracellular Anx A5 bound to PS, transiently exposed at the surface of osteoclasts at the time of fusion ^18^, facilitate La association with the surface of differentiating osteoclast precursors.

To summarize, while on its own La has no fusogenic activity, at the surface of differentiating osteoclasts it promotes fusion of their membranes. This promotion does not require La domain, RRM1 or the NLS of La and may involve complexes that La forms with Anx A5, one of the previously identified components of the osteoclast fusion machinery.

#### La presents a potential target for influencing osteoclast formation and function

The previously elucidated relationship between osteoclast fusion and bone resorption ^8–10^ led us to hypothesize that by regulating osteoclast size, La also regulates bone resorption. We evaluated this hypothesis by differentiating osteoclasts on fluoresceinated calcium phosphate, a biomimetic of bone, and assessed osteoclast-dependent bone resorption by the release of fluorescein into the media (illustrated in Fig. 5a). Monocyte derived precursors (only M-CSF) released minimal trapped fluorescein, but the addition of RANKL resulted in formation of multinucleated osteoclasts that readily resorbed calcium phosphate and released fluorescein (Fig. S1b). Overexpression of La 1-375 promoted bone resorption, while the uncleavable La mutant, D371,374A, had no effect (Fig. 5b). Moreover, RNAi mediated reduction of La reduced bone resorption by ~40% compared to non-targeted controls (Fig. 5c). a-La antibodies that inhibit fusion (Fig. 3e) also dramatically reduced osteoclast-dependent bone resorption in a dose dependent manner (Fig. 5d). Finally, the extracellular addition of recombinant La 1-375 to fusing osteoclasts dramatically increased osteoclast bone resorption (Fig. 5e). From these data, we conclude that targeting cell surface La bidirectionally regulates both osteoclast fusion and subsequent bone resorption.

**Figure 5:**
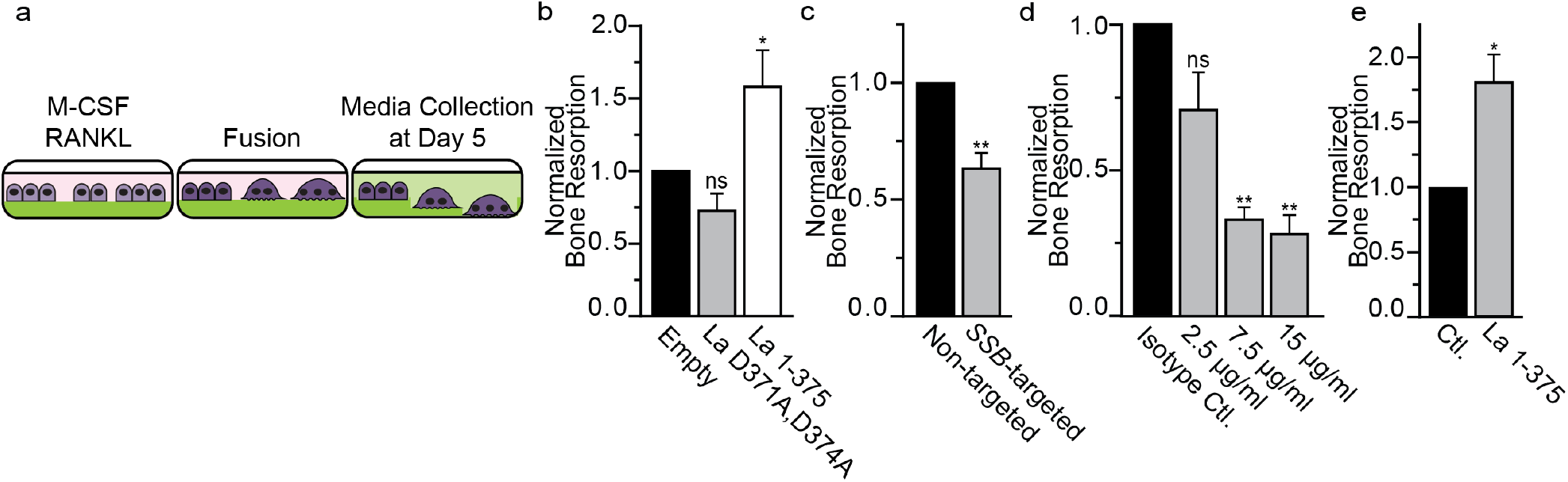
Interfering with La influences bone resorption by human osteoclasts. **(a)** Illustration depicting the use of fluoresceinamine-labeled chondroitin sulfate trapped in calcium phosphate coated plates to assay bone reportion. **(b)** Bone resorption in osteoclasts overexpressing uncleavable La (D371A, D374A) or truncated La (1-375) normalized to that for the osteoclasts transfected with empty plasmid. (n=3) (P=0.0564 and 0.0471) **(c)** Bone resorption in osteoclasts transfected with siRNA targeting La transcript normalized to that for non-targeted siRNA. (n=5) (P=0.0045) **(d)** Bone resorption in osteoclasts exposed to different concentrations of α-La normalized to that for the cells treated with isotype control antibodies at 7.5 μg/ml. (n=3) (P=0.0723, 0.0098 and 0.004) **(e)** Bone resorption in osteoclasts treated with 40nM recombinant La 1-375 normalized to the control after adding the same amount of PBS. (n=3) Statistical significance was assessed via paired t-test. Error bars = SEM.

In biologically relevant situations, osteoclastogenesis develops in the context of interactions between osteoclast precursors with bone-forming osteoblasts/osteocytes and other cell types, generating much lower concentrations of RANKL and many other osteoclastogenesis-regulating factors ^50^. To explore whether La is involved in osteoblast-induced osteoclast formation, we co-cultured primary human osteoblasts isolated from trabecular bone and human osteoclast precursors, derived via M-CSF induction of primary human monocytes. Osteoblasts and osteoclast precursors were cultured isolated from each other by well inserts (Fig. 6a). Without removing well inserts, we observed no fusion between osteoclast precursors. Upon removal of well inserts, media from the osteoblast/osteoclast wells mixed and cocultured osteoclast precursors rapidly fused to produce multinucleated osteoclasts. Addition of α-La antibody blocked nearly 75% of the fusion between osteoclasts in such co-cultures (Fig. 6b,c) confirming the involvement of La in osteoclast formation in a biologically relevant model of bone remodeling lesions.

**Figure 6:**
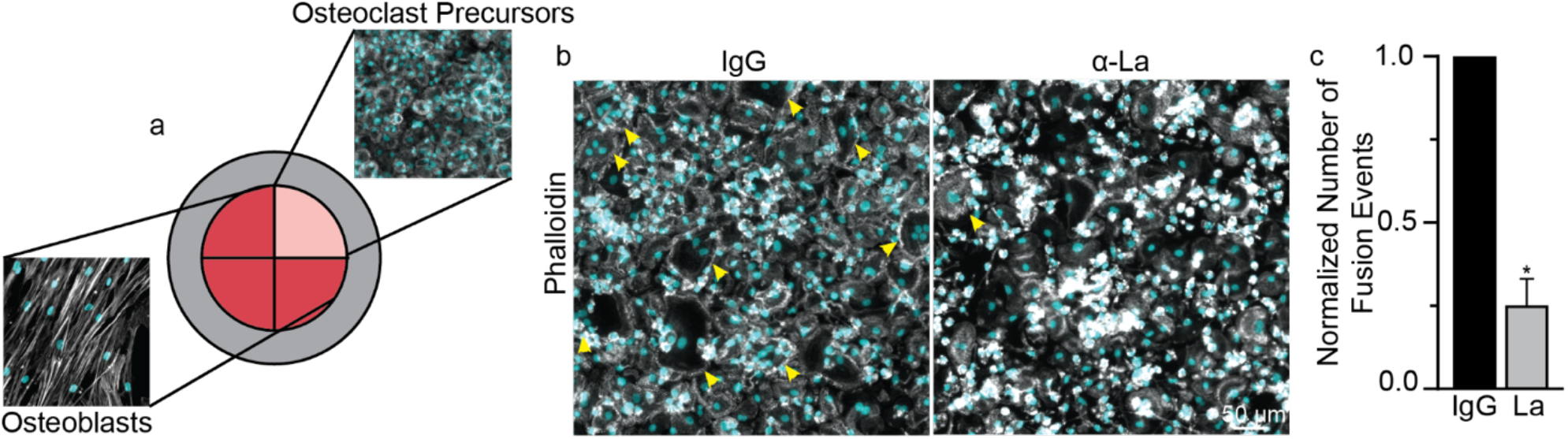
Osteoclast formation in human osteoblast/osteoclast precursor co-culture depends on La protein. **(a)** Schematic of the multi-well configuration of the human osteoblast/osteoclast co-culture system used. Multi-well dividers were removed following the M-CSF derivation of human monocytes, and osteoclast precursors and primary osteoblast media intermixed. (Grey=Phalloidin-Alexa488, Cyan=Hoechst) **(b)** Representative immunofluorescence images comparing osteoclast fusion with control IgG or α-La antibodies. Arrowheads denote syncytia with ≥3 nuclei. **(c)** Quantification of osteoclast fusion in the co-cultures in the presence of α-La normalized to that in the presence of control IgG. (n=4) (P=0.0331) Statistical significance was assessed via paired t-test. Error bars = SEM.

To explore whether La function plays a role in bone pathology, we have focused on fibrous dysplasia of bone (FD), an osteoclast-dependent bone disease ^51^. FD is caused by gain-of-function mutations in Gαs that lead to constitutively increased cAMP signaling and upregulation of cAMP/RANKL-dependent osteoclastogenesis ^52^. In a conditional, tetracycline inducible mouse model, FD-like bone lesions, develop in adult mice within 2 weeks following doxycycline (Doxy) administration ^53^. The formation of these lesions is driven by activation of an inducible gain-of-function mutant, Gα_s_^R201C^, specifically in cells of the skeletal stem cell linage responsible for the excessive RANKL production observed in FD. This excessive RANKL production results in the ectopic formation of numerous, large osteoclasts that excessively erode healthy bone. Using bone marrow explants from these FD mice, we established a novel, robust *ex vivo* model of the ectopic osteoclast formation observed in FD (Fig. 7a). As depicted in Fig. 7b, culture of these FD explants in the presence of M-CSF alone resulted in numerous adherent cells but no multinucleated, TRAP^+^ osteoclasts. In contrast, addition of Doxy resulted in the rapid development of fibrous cell clumps (arrow) and numerous multinucleated, TRAP^+^ osteoclasts (arrowheads) that were not observed in explants from wild-type littermates, lacking the inducible Gα_s_^R201C^ element. Doxy-induced osteoclastogenesis was accompanied by a ~17-fold increase in mRANKL produced by the explants (Fig. 7c). Importantly, α-La antibodies blocked osteoclast fusion elicited by the addition of Doxy to FD explants by ~60% and reduced the number of multinucleated osteoclasts observed by ~40% (Fig. 7d-f).

**Figure 7:**
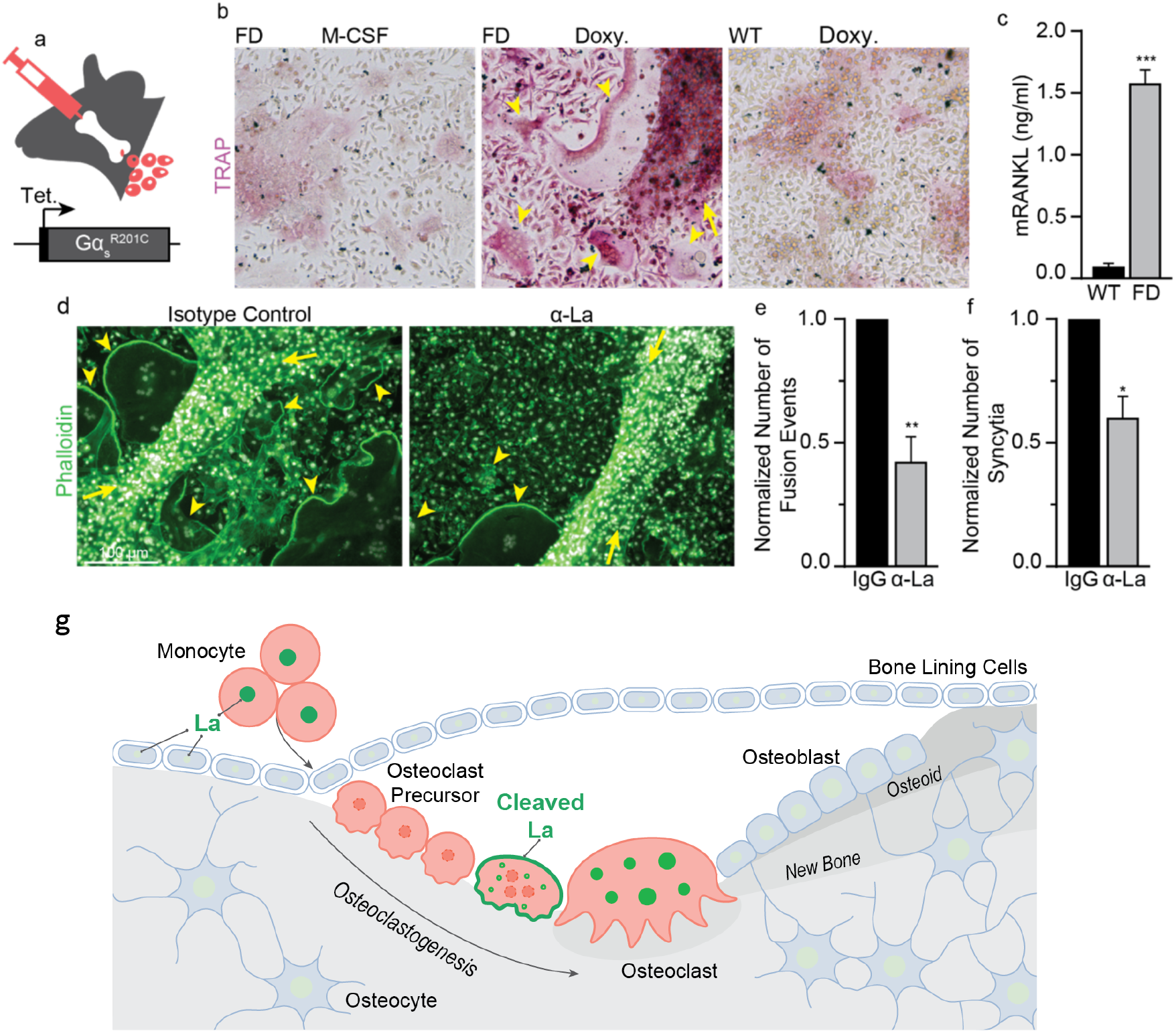
α-La treatment suppresses ectopic osteoclast formation in FD explants confirming that La regulates osteoclast activity in health and disease. **(a)** Illustration of *ex vivo* bone marrow culture system based on a tetracycline-inducible model of FD. **(b)** TRAP staining of bone marrow explants from a homozygous Gα_s_^R201C^ mouse (FD) and a wild-type littermate (WT). Explants were either cultured with M-CSF alone, or M-CSF and Doxy. **(c)** ELISA quantification of mRANKL produced in FD vs WT cultures following Doxy treatment. (n=4) (P=0.0005) **(d)** Representative images of FD explants activated by Doxy treated with isotype control or α-La antibody **(e)** Quantification of the number of fusions producing osteoclast syncytia with ≥3 nuclei. (n=4) (P=0.0054) **(f)** Quantification of the number of osteoclast syncytia with ≥3 nuclei from **d**. (n=4) (P=0.0178) In **b** and **d,** arrowheads denote multinucleated osteoclasts and arrows denote fibrous cell masses developed after Doxy addition and characteristic for FD. Statistical significance was assessed via unpaired t-test in **c** or paired, t-tests in **e** and **f.** Error bars = SEM. (**g**) An illustration of La protein in the process of osteoclast formation. La (green) carries out its canonical, ancient function in the nuclei of all eukaryotic cells as an essential, ubiquitous RNA binding protein. We propose that La has an additional, specialized function in the formation of multinucleated osteoclasts. In osteoclastogenesis, La dissipates as circulating monocytes become osteoclast precursor cells. When osteoclast commitment is initiated by RANKL, La returns but is quickly cleaved by proteases and shuttled to the surface of fusing osteoclasts. At the surface of fusing osteoclasts, La plays a novel function as a membrane fusion manager. When osteoclasts arrive at the “right size” for their biological function, surface La dissipates and is replaced by canonical, non-cleaved La that returns to the nuclei of mature osteoclasts mature osteoclasts.

To summarize, cell-surface La regulates formation of human and murine multinucleated osteoclasts triggered by biologically relevant interactions between osteoclast precursors and bone-forming cells. Targeting La modulates fusion between osteoclast precursors and, in turn, alters the propensity of the resulting osteoclasts to resorb bone. Furthermore, our findings that La is involved in osteoclast formation in an *ex vivo* FD model suggest that targeting La function at the surface of developing osteoclasts can be an effective therapeutic intervention in FD, and likely other resorptive bone diseases stemming from excessive osteoclast activity.

## DISCUSSION

### A novel function of La protein

Here we report that the differentiation of murine and human monocytes into multinucleated osteoclasts is dependent on tightly choreographed changes in the steady-state level, post-translational modification and cellular localization of La (Fig. 7g). These changes and our conclusion that cell surface-associated La regulates osteoclast fusion are startling in the context of the vast literature covering La’s role in RNA biology. La is generally thought of as an abundant, ubiquitous, mostly phosphorylated RNA-binding protein largely confined to the nucleus in virtually all eukaryotic cell types ^26,54^ However, at the onset of osteoclastogenesis, M-CSF-derived precursors show a dramatic loss of La protein, suggesting that this differentiation process may require the concerted downregulation of a specific La-regulated pool of mRNAs triggered by the loss of steady state La. In the following RANKL-induced stages of osteoclastogenesis, La reappears as a non-phosphorylated, proteolytically cleaved species in the cytoplasm and at the surface of the fusing osteoclast precursors. When the growth of osteoclasts slows, in the late stages of fusion, La is observed at its conventional size and nuclear localization. The rate of formation, the sizes of multinucleated syncytia and the subsequent bone resorption activity of osteoclasts are regulated by cell-surface La protein. In fact, cell-surface La regulates osteoclast functions by modulating the membrane fusion stage of osteoclast formation, not upstream differentiation processes. Lowering the amount of La by suppressing the steady-state level of its transcript, blocking the proteolytic processing required for its trafficking to the surface of the cells, or inhibiting its activity with antibodies inhibits fusion. Conversely, increasing La’s steady-state concentration by either overexpression or application of recombinant protein promotes fusion. In summary, these data demonstrate that La, a key protein in the RNA biology of eukaryotic cells, lives a second life at the surface of osteoclasts where it moonlights as a master regulator of osteoclast membrane fusion.

Our data demonstrate that La’s role in regulating osteoclast fusion and bone resorption is separate from the well-described canonical functions of La, and represents a novel function for the La protein. First, our ability to inhibit or promote synchronized osteoclast membrane fusion by non-membrane permeable reagents (e.g., antibodies, recombinant La) indicates that the regulation of osteoclast fusion depends on surface La. In contrast, well-characterized functions of La in the processing and metabolism of a variety of different RNAs ^26,27^ and in sorting microRNAs into extracellular vesicles ^55^ are carried out in the nucleus or cytoplasm and depend on La domain-RNA interactions. Only in some special biological processes, including in herpes simplex virus and adenovirus infections ^56,57^, adding serum to serum-starved cells ^32,58^, and in the early stages of apoptosis ^33,42^, is La protein found exposed at the surface of the viable cells ^33,42^. However, the only suggested function of these previous examples of cell surface bound La has been to recruit regulatory T cells to damaged tissues in an effort to downregulate an immune response during cell death ^59^.

In another striking distinction from the well-characterized functions of La in RNA metabolism, La regulation of osteoclast formation does not depend on interactions between the highly conserved La domain and RRM1 of La with RNA. This conclusion is supported by our finding that neither mutations in critical residues within the La domain nor deleting the entire N-terminal half of La protein (containing both the La domain and RRM1) abolishes the ability of the recombinant La to promote osteoclast fusion. Since recent studies found RNAs present on the surface of living cells that are involved in monocyte interactions ^60,61^, it remains possible that RNA is involved in surface La’s role in regulating osteoclast fusion. However, even if the function of La in osteoclast formation depends on La-RNA interactions at the cell surface through some yet unknown mechanism, this novel function fundamentally differs from the classical functions of La dependent on its La domain- and RRM1-mediated RNA binding in the nucleus and cytoplasm.

The mechanisms by which cell-surface La regulates osteoclast fusion remain to be clarified. Since La, on its own, initiates neither hemifusion nor fusion between bound membranes, it is unlikely that La directly catalyzes and/or drives membrane fusion. More likely La recruits or stimulates other components of the osteoclast fusion complex. The latter scenario is supported by our findings that highlight La’s association with the fusion regulator Anx A5. Anx A5 has been implicated in several cell-cell fusion processes (reviewed in ^49^). In the case of osteoclast fusion, osteoclastogenic differentiation of human monocytes is associated with a strong increase in the amount of Anx A5 present at the cell surface and treatments suppressing the expression and activity of cell surface Anx A5 inhibit synchronized osteoclast fusion ^18^. In contrast to Anx A5, two other members of the Annexin protein family: Anx A1, which is not involved in human osteoclast fusion ^18^, and Anx A4, which has many structural and functional similarities to AnxA5 ^62^, were not found in supramolecular complexes with La. The lack of Anxs A1 and A4 but the presence of Anx A5 in La protein complexes further support the hypothesis that La specifically associates with partners involved in the fusion stage of osteoclast formation. Specific mechanisms by which interactions between La and Anx A5, and possibly, La interactions with other components of osteoclast fusion machinery, regulate cell-cell fusion remain to be clarified. We find that recombinant La and Anx A5 directly interact, and that Anx A5 facilitates the association of La with membranes containing PS in a Ca^2+^-dependent manner. These observations in combination with the previously reported dependence of osteoclast fusion on cell surface PS and Anx A5 ^18^ begin to shed additional light on how osteoclasts may employ PS to trigger the assembly of a fusion complex between committed precursors.

While we found La to promote and regulate the formation of multinucleated osteoclasts, we do not know yet whether La is indispensable for osteoclast formation. La is required for mouse development and the establishment of embryonic stem cells ^38^, and efforts to specifically delete La in B cell progenitors and the forebrain have resulted in the loss of these cell types ^37^. Importantly, while the steady-state levels of La are quite low in osteoclasts precursors, these levels can be sufficient to support RNA metabolism. Futher resolution of the regions within La that are critical for its role in osteoclast fusion may afford us the ability to perturb La’s fusion function, while leaving its essential, RNA chaperone functions intact. This resolution will be essential for testing whether La is essential for osteoclast membrane fusion.

Our findings add La to a growing list of “moonlighting proteins” that serve several, sometimes strikingly unrelated, functions ^63,64^. For instance, the ubiquitous housekeeping proteins glycolytic enzyme Glucose-6-phosphate isomerase and RNA-binding protein nucleolin localized primarily in the cytoplasm and nucleolus, respectively, have second, unrelated functions at the cell surface ^63,65^. For La specifically, the N-terminal La motif is very highly conserved from yeast to man ^26^. However, latter domains of La, particularly the C-terminal half of the protein, are only weakly conserved. To this point, from yeast to vertebrata La experienced ~50% increase in its molecular mass due to an expansion in its C-terminus. Our data suggest that while the La motif, conserved throughout a billion years of evolution, is responsible for La’s original functions in RNA metabolism, regions in La’s C-terminal expansion, yet unidentified, carry out the protein’s more recently acquired function in regulating osteoclast formation.

The specific contributions of the different regions of La protein to its different functions; the evolutionary processes by which La protein has acquired its role in the formation of multinucleated osteoclasts; and the mechanisms that control the delivery of NLS-lacking La species to the surface of cells at the right time will have to be explored in future work. Intriguingly, La is not the only RNA-binding protein implicated in cell-cell fusion. Both Acheron, a member of La protein family ^66^, and HuR, a member of ELAV-family of RNA-binding proteins ^67^, are involved in formation of multinucleated myotubes. Comparison of the contributions of different RNA chaperones to diverse cell fusion processes will clarify whether cell fusion control by moonlighting RNA-binding proteins represents a conserved functional motif in the mechanisms human cells use to create syncytial cell types.

### La-dependent stage of osteoclastogenesis as a potential target for therapies

Imbalance of bone-formation and resorption in many skeletal diseases is linked to either excessive (e.g., osteoporosis, Paget’s disease and FD), or insufficient activity of osteoclasts (e.g., osteopetrosis). Here we report that the formation of multinucleated, human osteoclasts can be inhibited or promoted by treatments targeting La protein at the surface of osteoclast precursors. Importantly, α-La antibodies inhibit fusion and bone resorption by osteoclasts derived from RANKL activated monocytes. α-La antibody treatment also inhibited the formation of multinucleated osteoclasts in human osteoclast precursor/osteoblast co-cultures modeling bone remodeling lesions, where osteoclastogenic factors are produced by osteoblasts within the lesion ^68^. Moreover, our hypothesis that cell-surface La plays an important role in osteoclast formation within biologically relevant contexts is further substantiated by our experiments in a novel *ex vivo* model of FD. Development of FD is characterized by drastically increased levels of RANKL and other osteoclastogenic factors in serum and the excessive, ectopic formation of numerous multinucleated osteoclasts in the vicinity of bone lesions. As expected, induction of the FD phenotype in bone marrow explants resulted in high concentrations of RANKL and ectopic osteoclastogenesis. Finding that α-La inhibits the formation of multinucleated osteoclasts, both in size and number, confirms the importance of La as a novel target in bone pathologies and highlights the potential of La as a target for future therapeutic development.

Some of the proteins involved in the early stages of osteoclastogenic differentiation have already been tested in animal and/or clinical studies as potential therapeutic targets ^69^. The α-RANKL antibody denosumab is an FDA-approved drug for the treatment of osteoporosis ^21,70^. La-dependent osteoclast fusion, downstream of the RANKL/RANK/osteoprotegerin signaling pathway, presents a novel target for therapies at a different mechanistic stage of bone remodeling. Taking into account that mononucleated osteoclasts do resorb bones, blocking La-dependent osteoclast fusion is expected to have more subtle and selective effects on bone resorption than blocking the upstream formation of osteoclast precursors with α-RANKL antibodies ^71–73^. Like RANKL, cell surface La is accessible for cell-impermeable drugs. In some clinical situations, more subtle action of La-targeting treatments can be advantageous. Furthermore, unlike RANKL, which in addition to osteoclastogenesis regulates immune response ^74^, the only known function of cell surface La is its newly identified role in regulating osteoclast fusion. Surface La’s specificity may minimize off-target effects. Finally, osteoclasts are known to release factors that regulate osteoblast activity ^75^. Blocking osteoclastogenesis altogether by targeting RANKL likely blocks osteoclast-osteoblast signaling. Suppressing the fusion stage of osteoclast formation by targeting La, while maintaining the ablity of osteoclast to differentiate, may maintain this osteoclast-osteoblast crosstalk within the bone remodeling lesion.

In summary, our data demonstrate a function of La protein as a key regulator of osteoclast formation, a role strikingly different in place of action, mechanism, and partner protein (Anx A5) from its well-recognized functions as an RNA-chaperone. The future development of safe and effective reagents targeting cell-surface bound La may lead to novel antiresorptive therapies for osteoporosis, mechanistically orthogonal to the existing approaches. We expect our new *ex vivo* resorptive bone disease model utilizing FD cells, which condensed to days the time course of excessive osteoclast formation typical of disease, represents a powerful new tool for the study of osteoclastoegensis in health and disease states.

## Author Contributions

J.M.W., E.L., and L.V.C. designed osteoclast fusion experiments. J.M.W. and E.L. performed these experiments and analyzed the data. J.M.W. has designed, performed and analyzed all biochemical experiments. J.M.W., L.F.D.C, M.T.C. and LV.C. have designed bone marrow explant experiments and J.M.W. has performed them and analyzed the data. Liposome experiments were designed, carried out and analyized by K.M. and J.M.W.. Immunofluorescence experiments and experiments with hemagglutinin expressing cells have been designed, performed and analyzed by E.L. and J.M.W.. S.M and R.J.M. have contributed to the work an unpublished triple mutant of La, which S.M. has characterized and advised J.M.W. in the use of this mutant. R.J.M. and M.T.C. have contributed to the design of the work and interpretation of the data within the context of La protein biology and bone biology, respectively. J.M.W. and L.V.C. wrote the manuscript with assistance from all authors.

## Acknowledgements

We thank Drs Marjan Gucek, Yanling Yang and the proteomics core at the National Heart, Lung and Blood institue for their assistance in identifying La protein. We thank the National Institutes of Health Department of Transfusion Medicine for isolating the monocytes used in this study. The research in L.V.C.’ and R.J.M.’ laboratories was supported by the Intramural Research Program of the Eunice Kennedy Shriver National Institute of Child Health and Human Development, National Institutes of Health. The research in M.T.C.’ laboratory was supported by the Intramural Research Program of the National Institute of Dental and Craniofacial Research, National Institutes of Health.

## Supplementary Materials

### Materials and Methods

#### Reagents

Human M-CSF and RANKL and murine M-CSF and RANKL were purchased from Cell Sciences (catalogue #CRM146B; #CRR100B; # CRM735B and CRR101D, respectively). LPC (1-lauroyl-2-hydroxy-sn-glycero-3-phosphocholine, #855475); PC (1,2-dioleoyl-sn-glycero-3-phosphocholine, #850375C); PS (1,2-dioleoyl-sn-glycero-3-phospho-L-serine, #840035C); lissamine rhodamine phosphatidylethanolamine (1,2-dioleoyl-sn-glycero-3-phosphoethanolamine-N-(lissamine rhodamine B sulfonyl, #810150C) were purchased from Avanti Polar Lipids. Bone Resorption Assay Kits were purchased from Cosmo Bio Co. (Catalogue # CSR-BRA-24KIT) and used according to the manufacturer’s instructions. Hoechst 33342 and phalloidin-Alexa555 were purchased from Invitrogen (#H3570 and A30106, respectively). TRAP staining reagents were purchased from Cosmo Bio Co. (#PMC-AK04F-COS). The fluorescent lipid PKH26 (PKH26GL-1KT) and carboxyfluorescein, CF (5-(and-6)-Carboxyfluorescein, mixed isomers, #C368) were purchased from Sigma and Invitrogen, respectively.

#### Animals

All animal studies were carried out according to NIH-Intramural Animal Care and Use Committee (ACUC) of the National Institute of Dental and Craniofacial Research approved protocols (ASP #19-897), in compliance with the Guide for the Care and Use of Laboratory Animals. A mouse model of fibrous dysplasia with inducible expression of hyperactive Gα_s_^R201C^ in cells of the osteogenic lineage (*52*) was used to obtain bone marrow explants (described below). For this study we used 12-18 week old females.

#### Murine Bone Marrow Explant Culture

The tibia and femur were dissected from an inducible murine model of fibrous dysplasia described previously (*52*) or wild-type littermates. Holes were drilled into the epyphises of each bone using a 22-gauge hypodermic needle, and the bone marrow was flushed into a culture dish using alpha MEM. These bone marrow isolates were further dissociated through a fresh 22-gauge hypodermic needle to obtain a single cell suspension, and cultured in alpha MEM plus 20% FBS, 1x pen/strep and 1x Normocin (InvivoGen, # Ant-nr-1) for 7 days in T-75 culture flasks. Cells that adhered to the flask were washed 3 times with PBS and passaged using 0.05% Trypsin and a cell scraper and cultured for up to 3 passages in alpha MEM plus 20% FBS and 1x pen/strep. For Gα_s_^R201C^ expression induction in the bone marrow stromal cell subset of the explants, cells were plated at near confluency in 6-well plates and treated with 1 μM doxycycline (Sigma, # D9891-5G). During induction, media were refreshed daily. For antibody treatment, antibodies were added overnight when initial cell-cell fusion was observed (typically ~4 days of doxycycline treatment) at 6μg/ml overnight.

#### Cell cultures

##### Osteoclasts

Elutriated monocytes from healthy donors were obtained through the Department of Transfusion Medicine at National Institutes of Health. Cells were plated at ~ 2.9×10^5^ per cm^2^ in 35 mm dishes with polymer coverslip bottoms (Ibidi #81156) for imaging or 35 mm or 10 cm dishes for biochemical experiments in complete media [a-MEM supplemented with 10% Fetal Bovine Serum (FBS) and penicillin-streptomycin-L-glutamine (Gibco Invitrogene # 12571063; #26140079 and #10378016, respectively)]. Monocytes were differentiated to M2 macrophages in the presence of 100 ng/ml M-CSF for 6 days and then differentiated with 100 ng/ml M-CSF and 100 ng/ml RANKL for 3 days unless indicated otherwise. RAW 264.7 cells (ATCC, Manassas, VA, # TIB-71) were maintained in DMEM supplemented with 10% FBS to a maximum of 8 passages. RAW 264.7 cells were differentiated to osteoclasts in the presence of 100 ng/ml murine RANKL for 5 days. To separate unfused mononucleated and fused, multinucleated RAW 264.7 cells into separate fractions we have taken advantage of much stronger adherence of multinucleated cells to culture dish plastic. After washing in PBS, mixed RAW cultures (following RANKL differentiation) were left in Ca^2+^ and Mg^2+^ free PBS for 10 minutes at room temperature. Culture dishes were then tapped on the lab bench and a large portion of unfused, mononucleated RAW cells were released, and these released cells were collected via centrifugation. This process was repeated 2-4 times until dishes were left with a population of primarily fused, multinucleated syncytia. Mononucleated and multinucleated cell fractions were then processed for biochemical or imaging experiments as described below.

##### Human Osteoblast/Osteoclast Co-culture

Osteoblasts isolated from the trabecular bone of healthy individuals were obtained from PromoCell (#C12720) and cultured according to the manufacturer’s instructions. Osteoclast precursors were derived from primary human monocytes by 6 days of culture in M-CSF as described above. Osteoblasts and osteoclast precursors were cultured in 35mm dishes with 4-well culture inserts (ibidi, #81156) at a 3:1 well ratio. 48 hours before co-culture mixing, osteoblasts were switched to serum free alpha MEM with 1x pen/strep (Gibco). Following serum starvation, 4-well culture inserts were removed, and cells were cultured in their conditioned media overnight with or without treatment. Cells were fixed with 4% paraformaldehyde the following morning.

##### HAO-expressing cells and RBCs

NIH 3T3 mouse fibroblasts of clone 15 cell line that stably express influenza were a kind gift of Dr. Joshua Zimmerberg, NICHD, NIH (*76*). These HA0-expressing cells were cultured at 37°C and 5% CO_2_ in DMEM supplemented with 10% heat-inactivated FBS and antibiotics. The cells were used without trypsin pretreatment to keep HA in a fusion-incompetent form. Human red blood cells (RBCs) were isolated from the blood of healthy donors (who had consented to participate in the NIH IRB-approved Research Donor Program in Bethesda, MD; all samples were anonymized). RBC were labeled with the fluorescent membrane dye PKH26 and loaded with the fluorescent water-soluble dye CF, as described in (*77*).

#### Constructs & Recombinant proteins & Transfection

Recombinant La 1-408, 1-375, 1-375 Q20A_Y24A_D33I, 1-187 and 188-375 were amplified with primers designed with overlapping sequences and inserted between NdeI/HindIII in the V78 pET28A e. coli expression vector, adding to each a N-terminal 6xhis affinity tag. La 1-408 was expressed as a recombinant protein in E. coli and purified using IMAC columns by SD Biosciences (San Diego, CA). The remaining constructs were transformed into BL2 (DE3) chemically competent e. coli (Thermo Fisher Scientific) and protein expression was induced with IPTG. Cells were lysed with Bugbuster (Sigma), and 6xhis-La proteins were affinity purified using HisPur Cobalt Spin columns (Thermo Fisher Scientific), each according to the manufacturer’s instructions. Endotoxin contaminates were depleted from affinity purified 6xhis-La proteins using Pierce high-capacity endotoxin removal columns (Thermo Fisher Scientific), according to the manufacturer’s instructions. Proteins were then sterile filtered, aliquoted and kept at −80C.

Plasmids were introduced into primary osteoclasts at day 2 of RANKL stimulation via jetPRIME (Polyplus Transfection). FLAG-La 1-408, FLAG-La 1-375 and FLAG-La 1-375 Q20A_Y24A_D33I plasmids were a gift of the Maraia Lab (NICHD). Briefly, *SSB* (UniProt P05455) was inserted between HindIII and BamHI the pFLAG-CMV2 vector (Sigma). “Uncleavable” La was produced by taking the FLAG-La 1-408 plasmid and making two point mutations at amino acids D371A and D374A, abrogating the caspase cleavage sites at the C-terminal region of the protein (Emory Integrated Genomics Core). siRNA were introduced into primary osteoclasts after 1 day of RANKL stimulation via Lipofectamine RNAiMAX (Thermo Fisher Scientific). Non-targeted (Cat#4390843) and *SSB*-targeted (Cat#4392420_ID:s13469) siRNA were introduced at a concentration of 5ng/ml (Silencer Select, Ambion).

#### Antibodies

We used a-Cyclophilin B (CST, D1VdJ), a-GAPDH (CST, D16H11), a-Tubulin (Abcam, 7750), a-RANK (Abcam, 13918), a-Anx A5 (Abcam, 54775), control rabbit polyclonal IgG (Abcam, 27478), IgG2a (Abcam, 18415) used as an isotype control for a-La, Abcam, 75927), IgG1 (Abcam, 170190 used as an isotype control for a-Anx A5, Abcam, 54775), α-Anx A1 (Abcam, 47661), α-Anx A4 (Abcam, 65846), α-6xHis (Abcam, 18184) and a-FISH (Abcam, 118575).

The following anti-La antibodies were used for immunoblotting, immunostaining and immunoinhibition where indicated: Abcam, 75927 (referred to a-La); Anti-SSB antibody Invitrogen, PA5-29763 (referred to as a-LMW La); anti-La Phospho-Ser366 Abcam, 61800 (referred to as a-FL La). We characterized different a-La antibodies used in this study to identify the molecular species of La they recognized in human osteoclasts derived from monocytes (Table S1). To evaluate the selectivity of these antibodies for the two La species observed during osteoclastogenesis, we probed lysates from differentiating, human osteoclasts at the times when they primarily contained only one of the two La species described (i.e. mostly LMW La or mostly FL La). Abcam 75927 antibody (“a-La”) recognized both La species in Westerns. Invitrogen PA5-29763 (“a-LMW La”) preferentially recognizes LMW La. LMW La is a cleavage product of FL La; thus, FL La contains all the residues in LMW La. Our finding that a-LMW La preferentially recognizes LMW La suggests that this antibody must recognize a posttranslational or conformational epitope that differs in LMW vs FL La and not simply the primary amino acid sequence common to both. To specifically recognize FL La, we have used Abcam 61800 antibody to phosphorylated human La (phosphoSer366). While this a-FL La antibody did not work for Western blotting in our hands, in immunofluorescence staining, this antibody recognized FL La but not LMW La (see also (*33*)). In differentiating human osteoclasts at intermediate time point (Day 4 post RANKL addition), where Western blot analysis with a-La recognized both of osteoclast La molecular species, a-FL La exclusively recognized nuclear La, while a-LMW primarily recognized La in the cytoplasm.

#### Biochemical approaches

Cells were lysed on ice via pulse sonication and rotated end over end at 4°C for 45 minutes in the presence of protease inhibitors (cOmplete, Sigma, #118361700010). Steady-state protein levels were evaluated via SDS-PAGE followed by immunoblotting. Bulk proteins were evaluated via SDS-PAGE followed by silver stain (SilverQuest, Thermo Fisher Scientific). Bands of interest were cut from silver stained gels, destained and evaluated by liquid chromatography coupled with tandem mass spectrometry (Proteomics Core, NHLBI). The selective enrichment of cytosolic vs membrane associated protein fractions was carried out using Mem-PER™ Plus Membrane Protein Extraction Kit (Thermo Fisher Scientific, catalogue # 89842) according to the manufacturer’s instructions.

Immunoprecipitations were performed as described previously (*78*). Briefly, multi-protein complexes were sub-stocieometrically crosslinked using the non-membrane permeable, 12Å length, cleavable crosslinker 3,3’ Dithiobis(sulfosuccinimidylpropionate) (DTSSP) according to maufacture’s instructions (Thermo Fisher Scientific). Supermolecular complexes were immunoprecipitated using Sheep α-Ms IgG magnetic Dynabeads (Invitrogen) decorated with Ms antibodies targeting proteins of interest (α-La, Abcam, 75927; α-Anx A5, Abcam, 54775; IgG2a, Abcam, 18415; or IgG1, Abcam, 170190). Supramolecular complexes were denatured, crosslinking was cleaved via addition of reducing reagents (BME, BioRad), and proteins within these complexes were separated via PAGE. Proteins were transferred and probed for proteins of interest using immunoblotting (as described above). Rb antibodies were used to probe membranes for proteins of interest (α-La PA5-29763, α-Anx A5 14196, α-Anx A1 47661, or α-Anx A4 65846).

#### Transcript Analysis

For real-time PCR, total RNA was collected from cell lysates using PureLink RNA kit following the manufacturer’s instructions (Invitrogen # 12183018A). cDNA was generated from total RNA via reverse transcription reaction using a High-Capacity RNA-to-cDNA kit according to the manufacturer’s instructions (Applied Biosystems, # 4387406). cDNA was then amplified using the iQ SYBR Green Supermix (Biorad). All primers were predesigned KiCqStart SYBR Green primers with the highest rank score specific for the gene of interest or GAPDH control and were used according to the manufacturer’s instructions (Sigma). All Real-time PCR reactions were performed and analyzed on a CFX96 real-time system (Biorad), using GAPDH as an internal control. Fold-change of gene expression was determined using the ΔΔCt method (*79*). 3-4 independent experiments were performed and each was analyzed in duplicate.

#### Fusion Assays

Osteoclast fusion in our various culture systems was evaluated by fluorescence microscopy (*18*). Briefly, cells were fixed with 4% paraformaldehyde at timepoints of interest, permeabilized with 0.1% Triton X-100 and blocked with 5% FBS. Cells were then stained with phallodin-Alexa488 and Hoechst to label cells’ actin cytoskeleton and nuclei, respectively. 16 randomly selected fields of view were imaged using Alexa488, Hoechst and phase contrast compatible filter sets (BioTek) on a Lionheart FX microscope using a 10x/0.3 NA Plan Fluorite WD objective lense (BioTek) using Gen3.10 software (BioTek). Osteoclast fusion efficiency was evaluated as the number of fusion events between cells in these images, as described previously (*80*). Since regardless of the sequence of fusion events, the number of cell-to-cell fusion events required to generate syncytium with N nuclei is always equal to N-1, we calculated the fusion number index as ∑ (*N*_i_ – 1) = *N*_total_ - *N*_syn_, where *N_i_* = the number of nuclei in individual syncytia and *N_syn_* = the total number of syncytia. We normalized the number of fusion events to the total number of nuclei (including unfused cells) to control for small variations in cell density from dish to dish. In contrast to traditional fusion index measurements, this approach gives equal consideration to fusion between two mononucleated cells, one mononucleated cell and one multinucleated cell and two multinucleated cells. In traditional fusion index calculations, fusion between two multinucleated cells does not change the percentage of nuclei in syncytia. If instead one counts the number of syncytia, a fusion event between two multinucleated is not just missed but decreases the number of syncytia. In contrast, the fusion number index is inclusive of all fusion events.

#### Osteoclast Membrane Fusion Synchronization

Osteoclast fusion was synchronized as described in *(18, 81).* Briefly, osteoclast media was refreshed with 100 ng/ml M-CSF, 100 ng/ml RANKL and 350 μm lauroyl-LPC 72 hours post RANKL treatment. Following 16 hours, LPC was removed via 5 washes with fresh media and cells were allowed to fuse in the presence or absence of antibody treatment or recombinant La for 90 minutes.

#### HA0-RBC fusion assay

To test whether La is capable of mediating fusion, we applied this protein to HA0-expressing cells with RBCs tightly bound by the interactions between sialic acid receptors at the surface of RBCs and HA1 subunit of HA0 (*82*). HA0, an uncleaved form of HA, mediates binding but does not mediate fusion. HA-cells were twice washed with PBS and incubated for 10 min with a 1 ml suspension of RBCs (0.05% hematocrit). We washed HA-cells with zero to two bound RBC per cell with PBS to remove unbound RBC. Then the cells were exposed to 40 nM FL La.

Fusogenic activity (content mixing and/or lipid mixing) was assayed by fluorescence microscopy 1h after La application.

#### Liposome binding assay

Multilamellar liposomes were formed from pure PC or 9:1 (w/w) mixture of PC and PS. Both lipid compositions were supplemented with 0.5 mol % of lissamine rhodamine phosphatidylethanolamine. To prepare liposomes, lipid stock in benzene/methanol (95:5) was frozen in liquid nitrogen and freeze-dried overnight using SpeedVac (Savant). Dried lipid was resuspended in aqueous buffer (100 mM NaCl, 10 mM Hepes, pH 7.0) at 1 mM total lipid concentration and vortexed. Proteins and CaCl2 were added to the liposomes and the mixtures were incubated on ice for 30 minutes. To pellet, liposomes were centrifugated at 15,000g for 20 minutes to pellet, based on rhodamine fluorescence ~95% of liposomes. Centrifugated samples were then fractionated into a top, liposome-depleted fraction and a bottom fraction containing liposomes and liposome-bound proteins. Fractions were then solubilized via addition of Laemmli buffer (Bio Rad) and separated via SDS-PAGE, as described above. Recombinant La and recombinant Anx A5 were detected via their n-terminal 6xHis tag via α-6xHis antibody (Abcam) and signals for soluble vs liposome bound protein fractions were evaluated via densitometry. Data were presented as a percentage of protein signal bound to liposomes where

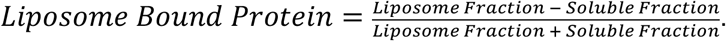

#### Fluorescence Microscopy Imaging

In the immunofluorescence experiments we washed the cells with PBS and fixed with warm freshly prepared 4% formaldehyde in PBS (Sigma, F1268) at 37^0^C. The cells were washed three times with PBS. To permeabilize the cells, we incubated them for 5 min in 0.1% Triton X100 in PBS. The cells were again washed three times with PBS and placed into PBS with 10% FBS, for 10 min at the room temperature to suppress non-specific binding. Then the cells were incubated with primary antibodies for 1 hour in PBS with 10% FBS. After 5 washes in PBS, we placed the cells for 1h at room temperature in PBS with 10% FBS with secondary antibodies (either Anti-rabbit IgG Fab2 Alexa Fluor or Anti-mouse IgG Fab2 Alexa Flour ® 488, both Cell Signaling Technology, Catalogue # 647 4414S and # 4408S, respectively, in 1:500 dilution) and then again washed the cells 5 times with PBS.

In the experiments that required immunostaining of non-permeabilized cells (Fig. 3b,c), we first fixed the cells as above and then incubated them with primary antibodies (10μg/ml) for 10 min at 37^0^C. After 2 washes with the full medium and 3 washes with PBS, cells were fixed, as described above. After fixing, we washed the cells 3 times with PBS and placed them into PBS with 10% FBS, for 10 min at the room temperature to suppress non-specific binding. Then the cells were incubated with secondary antibodies, as described above (1 hour in PBS with 10% FBS at room temperature) and, finally, washed 5 times with PBS.

Images were captured on a Zeiss LSM 800 airyscan, confocal microscope using a C-Apochromat 63x/1.2 water immersion objective.

#### Bone Resorption

Bone resorption was evaluated using bone resorption assay kits from Cosmo Bio USA according to the manufacturer’s instructions. In short, fluoresceinamine-labeled chondroitin sulfate was used to label 24-well, calcium phosphate-coated plates. Human, monocyte-derived osteoclasts were differentiated as described above, using alpha MEM without phenol red. Media were collected at 4-5 days post RANKL addition, and fluorescence intensity within the media was evaluated as recommended by the manufacture.

#### Statistical analysis

Each graph presents data from three separate biological replicates repeated on independent occasions unless stated otherwise in the legend. Data were assembled and analyzed using GraphPad Prism 8.0. For each experiment, cells from the same passage, donor or animal were paired across the differing conditions described. All functional dependencies reported were observed in each independent experiment. However, as known for the human monocyte-derived osteoclasts ^10^, times course of osteoclastogenic differentiation and baseline extents of fusion considerably varied for monocytes from different donors. We analyzed statistical significance using a paired ratio t test, where raw values are logarithmically transformed and then assessed.

In our analysis of the HA0-RBC experiments, Wilson method based confidence limits for binominal proportion were calculated in R (v. 4.1.1) using binconf function of Hmisc package (v. 4.5.0).

---

**Figure S1:**
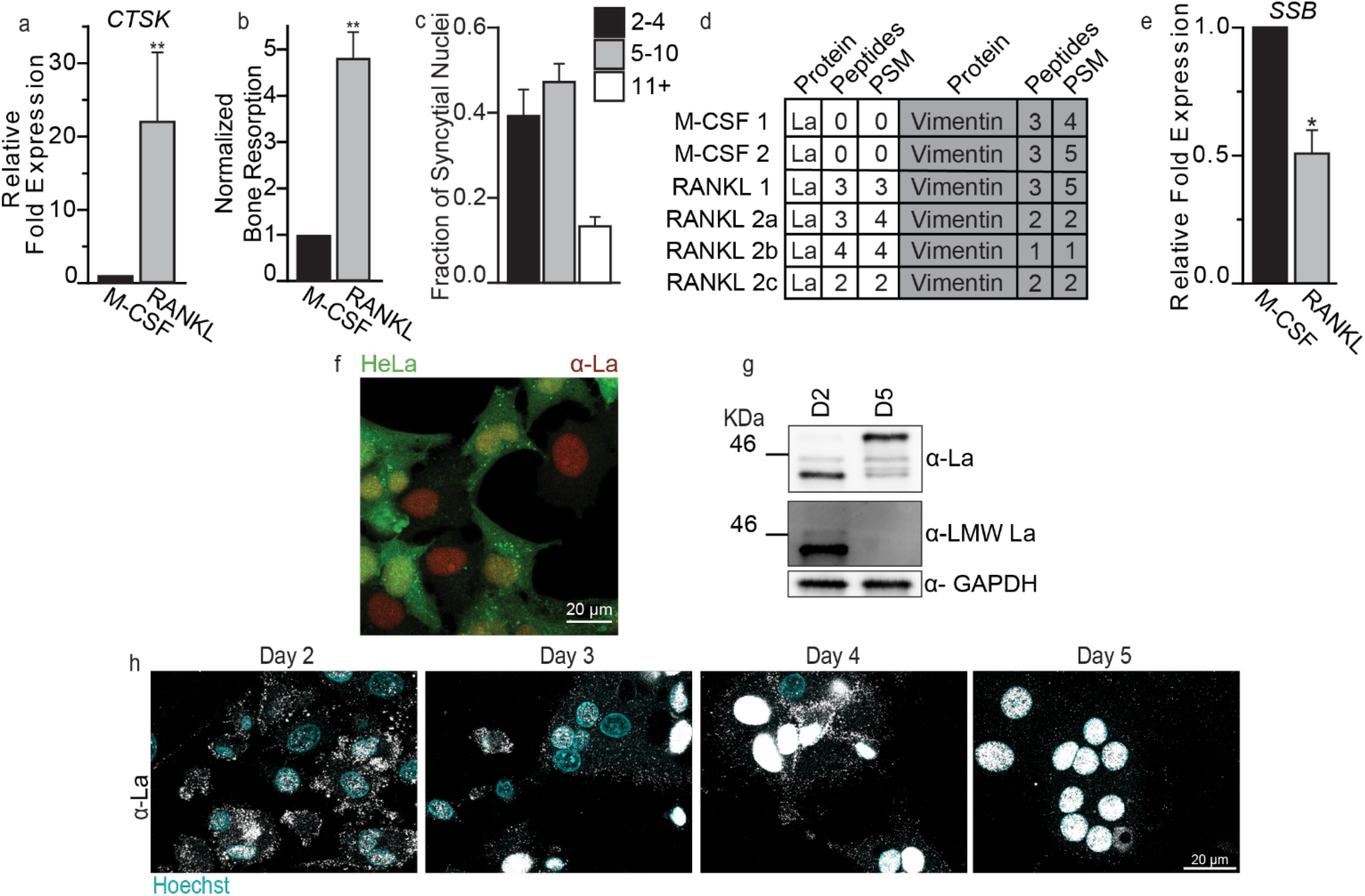
Characterization of monocyte derived osteoclasts and identification of La protein in osteoclasts. **(a)** qPCR evaluation of the osteoclast differentiation marker *CTSK* from the conditions depicted in Figure 1a. (n=4) (P=0.0079) **(b)** Quantification of fluorescent bone resorption in osteoclast precursors (M-CSF) vs differentiated osteoclasts (RANKL). (n=3) (P=0.0066) (Day 5 post-RANKL) **(c)** Fraction of nuclei in syncytial osteoclasts of varying sizes. (n=4) (Day 5 post-RANKL) **(d)** Mass spectrometry data from six excised bands separated in six separate lanes ran on a single gel, as seen in Figure 1b. Each lane represented a distinct cell lysate. Cells were collected from two healthy donors (1&2) and differentiated for six days with M-CSF and further differentiated for three days with M-CSF or M-CSF + RANKL. Cells from donor 2 were differentiated in three separate technical replicates (2a-2c). Vimentin was also detected in each sample, as it has a similar molecular weight to La, but vimentin levels were roughly equivalent in M-CSF vs RANKL samples. The table shows the total number of distinct peptide sequences identified in the protein group (Peptides) and the peptide spectrum matches (PSM). **(e)** qPCR evaluation of SSB (La gene) from the osteoclastogenic stages depicted in **a** and **d** (n=3) (P=0.027). **(f)** Representative immunofluorescence image of α-La staining in GFP expressing HeLa cells. **(g)** Representative tris-glycine Western blot evaluating La species predominate at 2 vs 5 days post RANKL addition. **(h)** Representative immunofluorescence images of La in forming osteoclasts 2-5 days post RANKL application (α-La, Abcam). Cells were stained for La at the described timepoints with membrane permeabilization. Statistical significance was evaluated via paired t-test. ** = P<0.001 Error bars = SEM.

**Figure S2:**
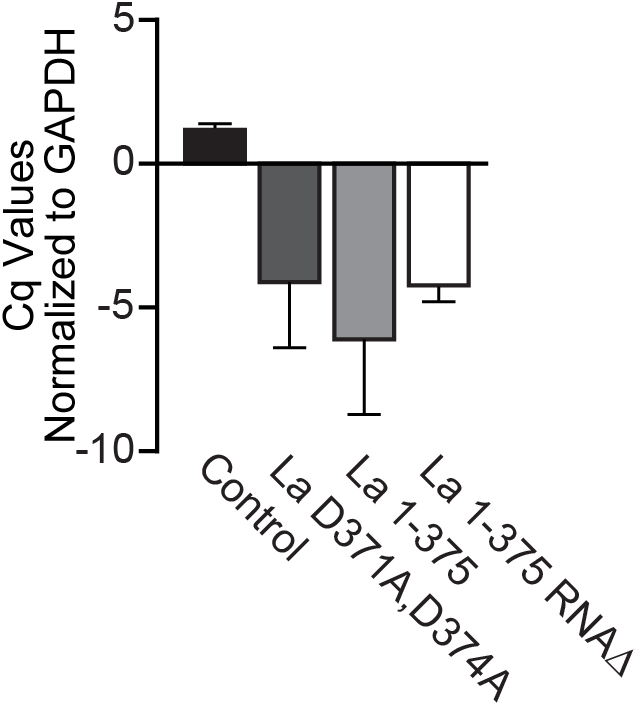
Exogenous La constructs express at similar levels in human osteoclasts. ΔCq values of La signal in human osteoclasts transfected with empty, La D371A,D374A, La 1-375 and “RNAΔ” La 1-375 Q20A_Y24A_D33I mammalian expression plasmids. GAPDH was used as a housekeeping transcript control. (n=2) Error bars = SEM.

**Figure S3:**
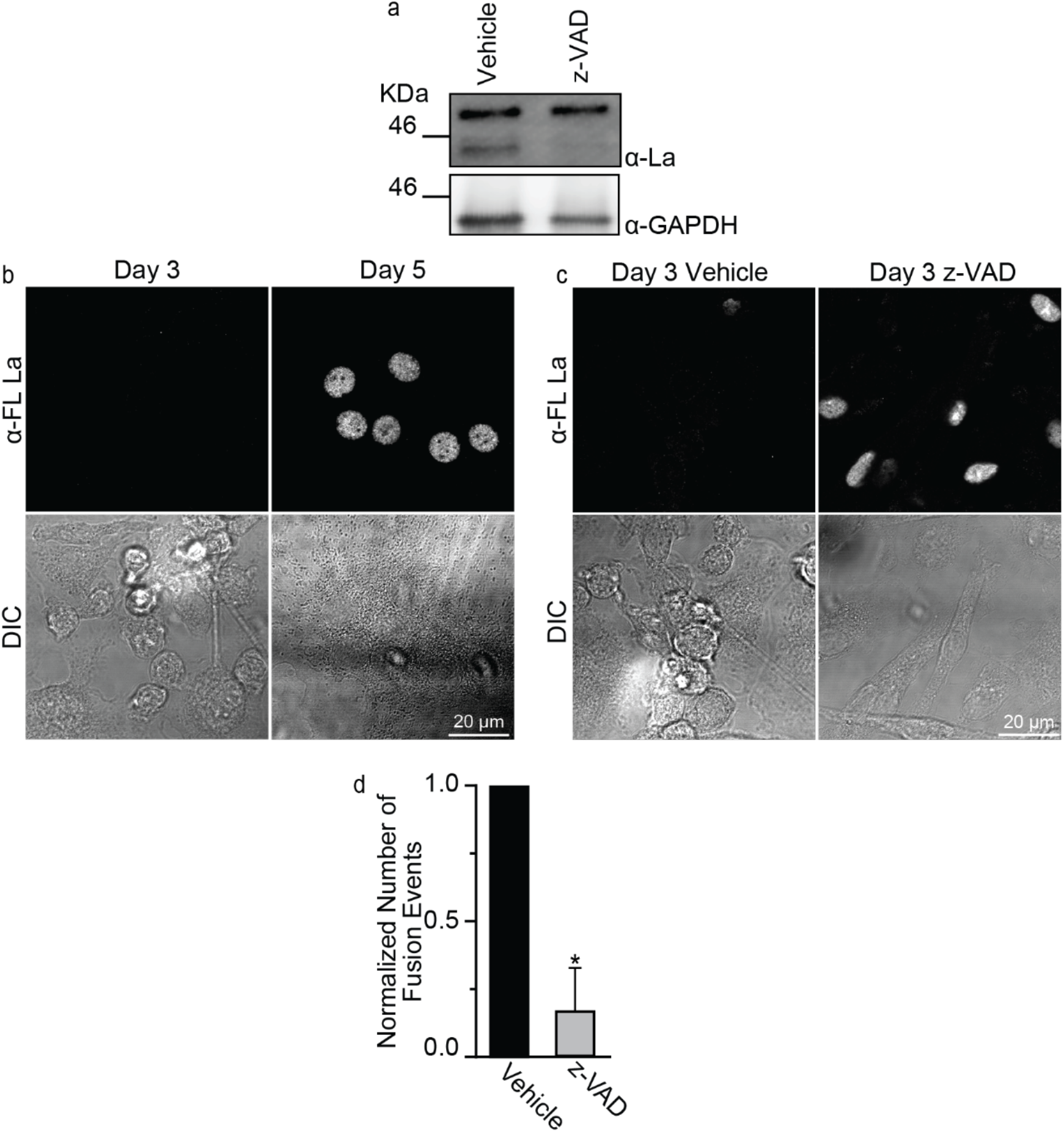
Formation of multinucleated osteoclasts depends on La cleavage. **(a)** Western blot demonstrating the ability of the caspase inhibitor z-VAD-fmk to block the production of cleaved La in human osteoclasts (Day 3-4 post-RANKL). **(b)** Representative immunofluorescence images of α-FL La staining in primary human osteoclasts during active fusion (Day 3) and when fusion is mostly complete (Day 5). **(c)** Representative immunofluorescence images of α-FL La staining in primary human osteoclasts during active fusion (Day 3) under control conditions (vehicle) and following inhibition of La cleavage via application of pan caspase inhibitor z-VAD-fmk. **(d)** Quantification of the number of fusion events in cells with 3+ nuclei from **c**. (n=3) (P=0.0455). Statistical significance was evaluated via paired t test. * = P<0.05. Error bars = SEM.

**Figure S4:**
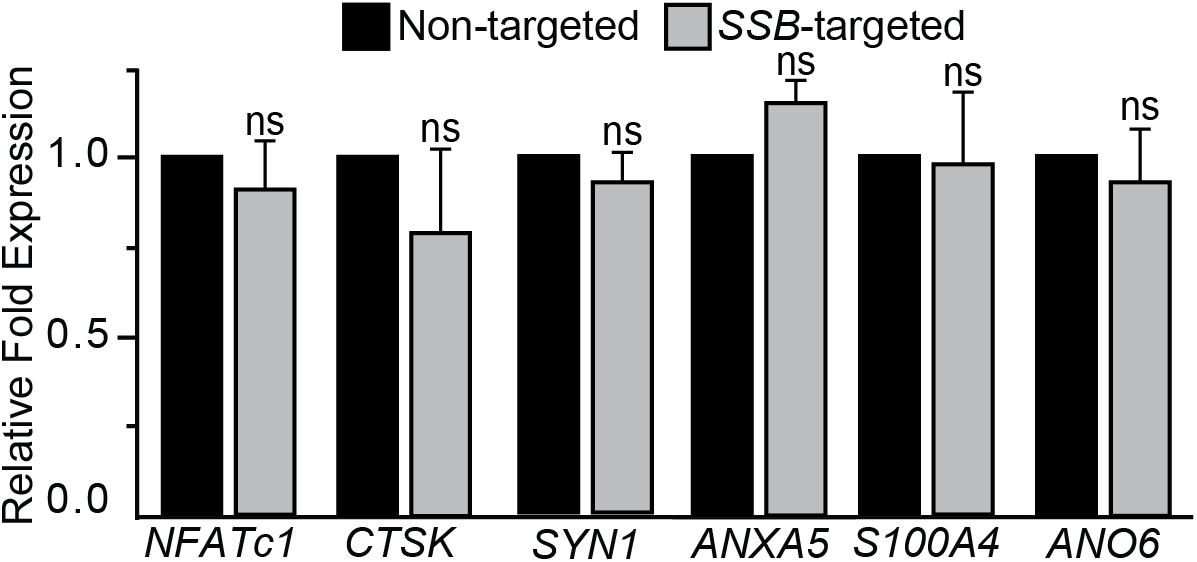
RNAi suppression of La does not alter the steady state transcript levels of proteins implicated in osteoclastogenic differentiation or osteoclast fusion. qPCR evaluation of two osteoclast differentiation markers, NFATc1 and CTSK and the osteoclast fusion related transcripts SYN1, ANXA5, S100A4 and ANO6 following the siRNA treatments described in Figure 2a. (n=4) Statistical significance was assessed via paired t-test. (P=0.4051, 0.4679, 0.45650, 0.6172, 0.7899 and 0.1129). Error bars = SEM.

**Figure S5:**
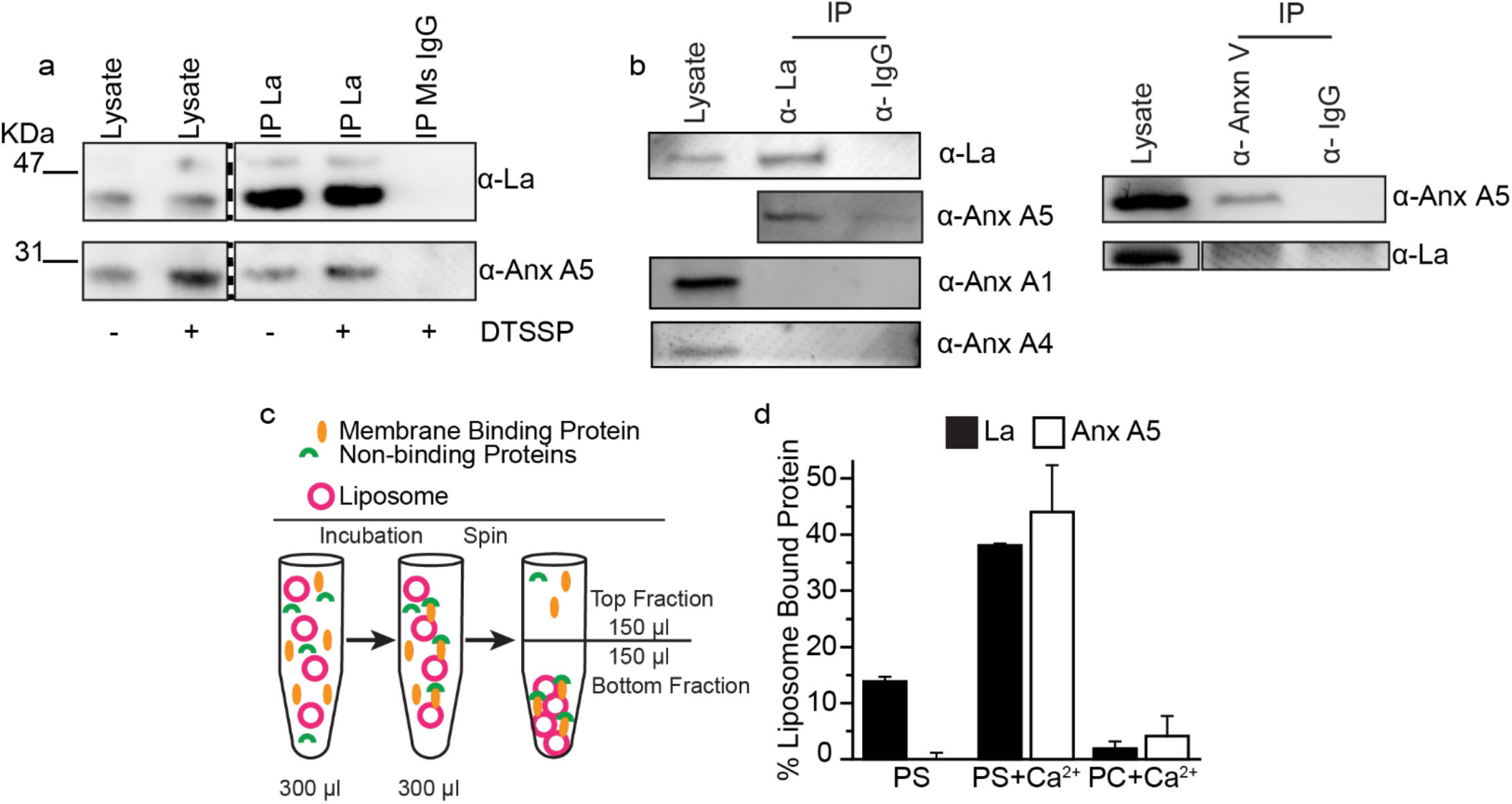
La interacts with Anx A5. **(a)** Immunoprecipitation of osteoclast lysates 3 days post-RANKL addition with or without DTSSP surface crosslinking. La supermolecular complexes were captured on immunomagnetic beads via mouse α-La or isotype control and complexes were blotted with rabbit antibodies raised towards the targets of interest. **(b)** Immunoprecipitation of osteoclast lysates with DTSSP surface crosslinking. La (left) or Anx A5 (right) supermolecular complexes were captured on immunomagnetic beads via mouse α-La or α-Anx A5, and complexes were blotted using rabbit antibodies raised towards targets of interest. **(c)** A cartoon illustration of our approach to identify membrane affinity by separating liposome bound proteins (Bottom Fraction) from soluble proteins (Top Fraction). **(d)** Quantification of recombinant La vs Anx A5 in liposome bound fractions, as illustrated in c. (n=2) Error bars = SEM.

**Table S1:** Antibodies used for the detection of La molecular species in human osteoclasts.

| **Antibody Producer/#** | **Referred to as** | **La Species Recognized in Human Osteoclasts** |
| --- | --- | --- |
| **Abcam/75927** | α-La | LMW La and FL La |
| **Invitrogen/PA5-29763** | α-LMW La | LMW La |
| **Abcam/61800** | α-FL La | PhosphoSer366 FL La |

